# A special envelope separates extra-chromosomal from mammalian chromosomal DNA in the cytoplasm

**DOI:** 10.1101/2023.03.02.530628

**Authors:** Laura Schenkel, Xuan Wang, Nhung Le, Michael Burger, Ruth Kroschewski

## Abstract

Expression from transfected plasmid DNA is generally transient, but little do we know on what limits this. Live-cell imaging revealed that DNA transfected into mammalian cells was either captured directly in the cytoplasm, or was soon expelled from the nucleus, upon its entry. In the cytoplasm, plasmid DNA was rapidly surrounded by a double membrane and frequently colocalized with extra-chromosomal DNA of telomeric origin, also expelled from the nucleus. Therefore, we termed this long-term maintained structure exclusome. The exclusome envelope contains endoplasmic reticulum proteins, the inner-nuclear membrane proteins Lap2β and Emerin but differs from the nuclear envelope by the absence of the Lamin B Receptor, nuclear pore complexes and by the presence of fenestrations. Further, Emerin affects the frequency of cells with exclusomes. Thus, cells wrap chromosomes and extra-chromosomal DNA into similar yet distinct envelopes. Thereby, they distinguish, sort, cluster, package, and keep extra-chromosomal DNA in the exclusome but chromosomal DNA in the nucleus, where transcription occurs.

## Introduction

In all eukaryotes the genome is enclosed in the nucleus, which compartmentalizes the chromosomes away from the cytoplasm (Güttinger et al., 2009). The separation between nucleoplasm and cytoplasm is ensured by a flat double membrane derived from the endoplasmic reticulum (ER). Exchange between the nucleoplasm and the cytoplasm occurs mainly through pores in this double membrane which are made selective by nuclear pore complexes (NPCs). Such NPC-containing double membrane is further specialized, e.g., by the presence of inner-nuclear membrane (INM) proteins, constituting the nuclear envelope. In many species, the nuclear envelope breaks down in mitosis and reassembles around the chromosomes upon mitotic exit. Whether nuclear envelope assembly is somehow restricted to chromosomes at the end of mitosis or if it can take place around any DNA in the cytoplasm and even throughout the cell cycle is unknown.

How mammalian cells assemble the nuclear envelope at the end of mitosis has been intensively studied. When the separated chromosomes are pulled to opposite spindle poles towards the end of anaphase, tubular ER membranes approach each segregating chromosomal mass from several sides establishing the beginnings of two nuclear envelopes (Anderson and Hetzer, 2007; Anderson and Hetzer, 2008). But still the trigger for membranes, being not necessarily exclusively ER, to approach and then contact the separated chromosomes is unresolved (Kutay et al., 2021; Schellhaus et al., 2016). Barrier-to-autointegration factor (BAF, BANF), which sequence unspecifically binds DNA, accumulates at the surface of mitotic chromosomes (Samwer et al., 2017; Zheng et al., 2000). BAF is required for wrapping the chromosomes in membranes, which contain initially several homogeneously distributed transmembrane proteins of the LAP2, Emerin, MAN1 (LEM)-domain family (Haraguchi et al., 2000; Haraguchi et al., 2001; Kobayashi et al., 2015). BAF binds to the LEM-domain of e.g., Emerin, and thereby establishes a DNA membrane tether (Lee et al., 2001). NPC assembly occurs after membrane patches established contact with the chromosomes and contribute to nuclear envelope sealing (Kutay et al., 2021; Otsuka et al., 2018). Would these events indistinctively take place and wrap up any type of DNA or are they exclusive to chromosomes?

In addition to the special case of mitochondria, there are situations *in vivo* where DNA is enwrapped by a membrane. For example, late in anaphase, lagging chromosomes can be enwrapped in an envelope and thus separate from the main nucleus, forming structures called micronuclei. Initially, the micronuclear envelope has all the characteristics of a nuclear envelope (Hatch et al., 2013). However, it degenerates over time. Over this period the enclosed DNA becomes fragmented and during one of the following mitoses the fragments reintegrate into chromosomes (Crasta et al., 2012; Zhang et al., 2015). In micronuclei, loss of Lamin B1 correlates with their decay (Hatch et al., 2013). It is not known, however, what makes micronuclei degenerate while the nucleus stays intact. Remarkably, lagging chromosomes were shown to be frequent. Yet, they were transient, hardly forming micronuclei, as they mostly reintegrated during the mitosis in which they appeared into the reforming nucleus in human non-transformed and transformed cell lines (Orr et al., 2021). Also in *Drosophila melanogaster* neuronal stem cells, chromatin fragments originating from chromosome ends were rarely found in micronuclei (Karg et al., 2015). Instead, these fragments either rejoined early in anaphase a membrane-free chromosomal mass or they rejoined newly formed nuclei via nuclear envelope channels and by tethering to chromosome ends later in anaphase (Karg et al., 2015; Warecki et al., 2020). Thus, lagging chromosomes mostly rejoin nuclei within the same mitosis or they form less frequently unstable micronuclei, where the fragmented micronuclear DNA ends up in nuclear chromosomes after some divisions. Remarkably, syncytia, which are cells with multiple stable nuclei, exist. All these nuclei are roughly of similar size as seen in the slime mold *Physarum polycephalum* and human osteoclasts (Gerber et al., 2022; Kopesky et al., 2014). In contrast micronuclei in mammalian cells are at least 5 times smaller than their corresponding nuclei (Kneissig et al., 2019). Therefore, separate nuclei and nucleus-like structures can form in the same cell but in many instances the smaller structures are unstable.

Remarkably, circular extra-chromosomal DNA excised from chromosomes, thus of endogenous origin, exists in every cell type tested (Noer et al., 2022; Paulsen et al., 2018). However, extra-chromosomal DNA can also be of exogenous origin. Remarkably, when lambda phage or plasmid DNA was mixed with *Xenopus* oocyte extracts, it was subsequently enwrapped by a nuclear envelope, which could suggest that any DNA can be enveloped by a nuclear envelope (Blow & Laskey, 1986; Newport, 1987). Thus, exogenous extra-chromosomal DNA introduction into cells by e.g., viral or bacterial infections or transfection provide opportunities to study the formation of nucleus-like structures, as well as help to understand how cells distinguish chromosomal from extra-chromosomal DNA. Therefore, in this study we have introduced DNA into the cytoplasm of mammalian somatic cells and characterized its enwrapping by membranes. Particularly, we investigated the similarities and differences between such membranes and a bona fide nuclear envelope.

## Results

### Cells sort transfected plasmid DNA to the cytoplasm

We chose to study the association of plasmid DNA with membranes after its transfection into human tissue culture cells, as its visualization is established. To visualize transfected plasmid DNA, we used the LacO/LacI-system, where a plasmid containing 256 repeats of the Lactose (Lac) Operon (pLacO) is introduced into cells stably expressing Lac Inhibitor (LacI) fusion protein with either GFP or mCherry as fluorescence tag. Here, we employed three different HeLa cell lines (hereafter referred to as “HeLa-LacI”) and transfected them with pLacO to analyze the localization of transfected plasmid DNA. Fluorescent LacI foci in the cytoplasm were detected in cells transfected with plasmid DNA either by lipofection or electroporation (two methods to introduce plasmid) (SFig. 1; as previously reported in (Wang et al., 2016)). Due to the presence of a nuclear localization sequence (NLS) in the LacI fusion protein, the nuclear LacI fluorescence generally masked signals coming from plasmids in the nucleus. These results confirm that the LacI foci report plasmid localization in cells after transfection; hereafter, we refer to these LacI foci as plasmid foci. We first used this reporter system to study the localization and dynamics of plasmid foci. To do so we performed time-lapse live-cell microscopy for up to 24 hours (Fig. 1A-D, SFig. 2, SFig. 3). We started image acquisition concomitantly with the addition of the plasmid-lipofection mix to the cells and under conditions that preserved viability in the dividing cell population (SFig. 2A).

**Fig. 1.**
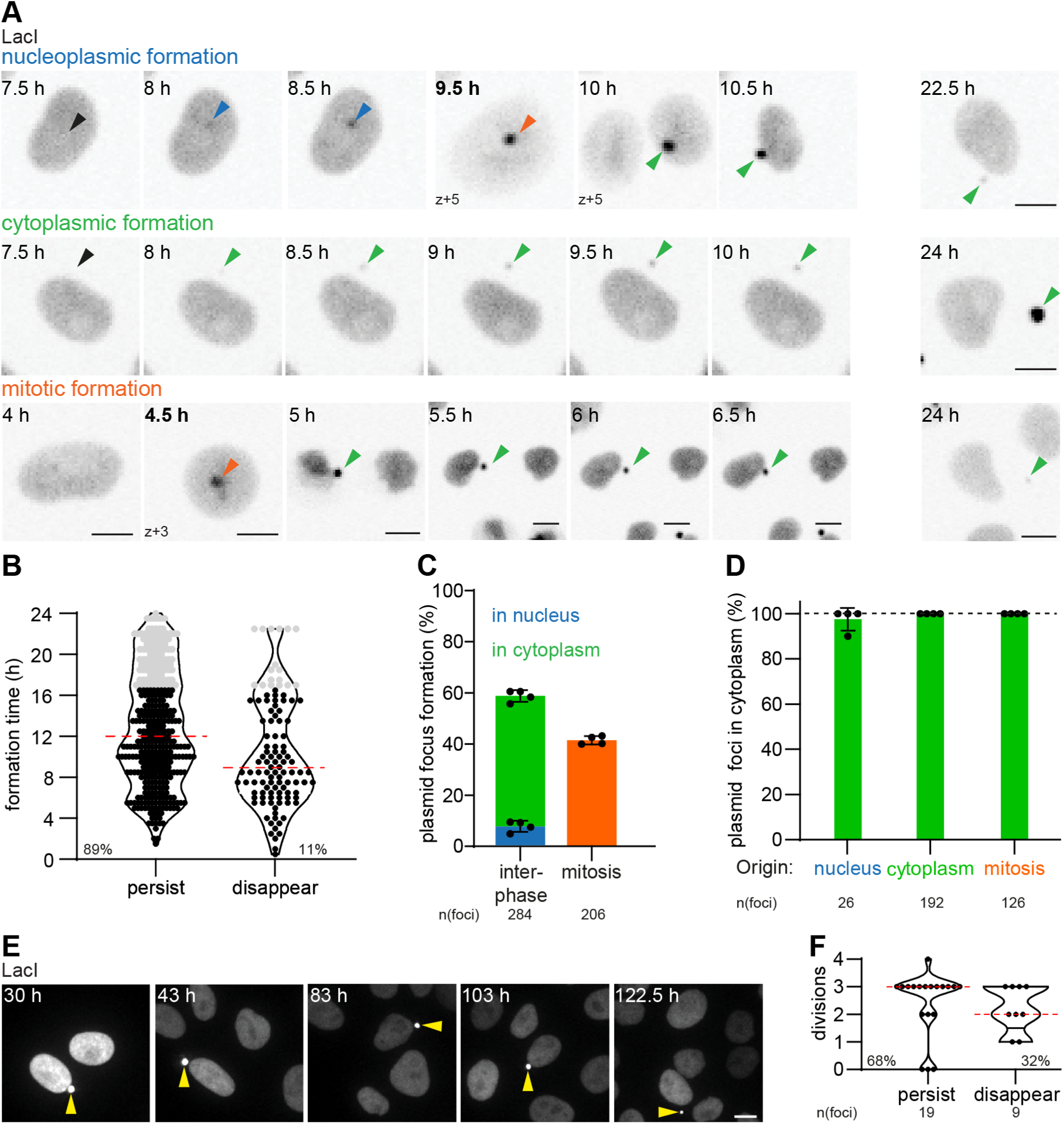
Cytoplasmic plasmid foci have 3 origins and are maintained in the cytoplasm long-term. (**A**) Time-lapse images of focus formations in HeLa-LacI cells lipofected at imaging start with pLacO. Scale bar, 10 µm. Time, after lipofection; bold time, mitosis. Arrowheads: nucleoplasmic focus, blue; cytoplasmic, green; mitosis, orange; future focus formation, black. Single z-slices. (**B**) Timing of individual focus formation events (circle) after pLacO lipofection. Persisting (persist) and disappearing (disappear) foci. 4 experiments (exp.) pooled; n(foci): 490; median, red. Last 25 % of all appearances, grey. % relative to all foci formed. (**C**) Focus formations in interphase or mitotic cells relative to all focus formations. 1 exp., circle; mean & SD. (**D**) Foci that are in the cytoplasm at imaging end depending on their origin. Last 25 % formations (in B) excluded. Color code as in (A). n(foci): 344; 100 % reference, dashed line. (**E, F**) Imaging started 30 hours after pLacO lipofection. 1 exp., n(cells): 28. (**E**) Time-lapse images of a pLacO transfected cell with one persisting focus, yellow arrowhead. Images, maximum intensity (max.) projected; (**F**) Maximal number of divisions a focus was detectable. One focus, circle. Persisting (persist) until imaging end or disappearing before (disappear); % relative to all foci.

We observed the formation of plasmid foci throughout the imaging period, perhaps because plasmids continuously entered the cells (Fig. 1B, SFig. 2B). Most of these plasmid foci (89 %) persisted throughout the entire imaging period (Fig. 1B). The other plasmid foci (11 %) were visible for variable durations (between 30 min and 17 hours) before disappearing (Fig. 1B, SFig. 2C). Consistent with our previous study, most cells exhibited only one plasmid focus (63 %; SFig. 2D, (Wang et al., 2016)). Next, we analyzed the history of cells with a single focus at the end of imaging (211 cells, 63 % of all cells with ≥ 1 plasmid foci, 19% of all imaged cells at the end of imaging). Interestingly, we found that 76 % of cells with one plasmid focus had either formed only a single plasmid focus or inherited it during mitosis. In contrast, partitioning of multiple foci during mitosis or disappearance of foci (21 % and 3 %, respectively) contributed less to the cells with one plasmid focus at the end of imaging. Notably, we did not observe any plasmid foci fusion events under our imaging conditions (SFig. 2E, F; total 291 plasmid foci in 114 cells over up to 24 hours). These data show that transfected cells usually form only one plasmid focus. Once formed, plasmid foci are generally stable.

Our live-cell imaging also revealed that 58 % of plasmid foci formed during interphase, while the other 42 % were plasmid foci formed during mitosis, away from the chromosomal mass (Fig. 1A, mitotic formation, bold time point, C, SFig. 2I). Amongst the plasmid foci formed during interphase 88 % formed in the cytoplasm and 12 % in the nucleus (Fig. 1A, C, SFig. 2G). Next, we analyzed the location of each appearing plasmid focus at the end of imaging. Irrespective of where and when plasmid foci formed, all but one ended up in the cytoplasm (Fig. 1A at 24 h, D, SFig. 2H, I).

Next, we wondered how plasmid foci formed in the nucleus entered the cytoplasm. Focusing on plasmid foci formed in the inner nucleoplasm, we observed two different translocation modes: 13 out of 15 such plasmid foci entered the cytoplasm during mitosis by being sorted away from the chromosomal mass (SFig. 3A (1^st^ example), B). The other two plasmid foci left the nucleus during interphase by nuclear budding (SFig. 3A (2^nd^ example), B), revealing that the cell employs at least two ways to exclude plasmid DNA from the nucleus.

These data make four points. First, most of the plasmid DNA becomes LacI-associated in the cytoplasm and remains there, possibly without ever reaching the nucleus. Second, there are two modes for nuclear plasmid to partition away from the chromosomes: through expulsion via nuclear budding from the interphase nucleus (SFig. 3A (2^nd^ example)) or through separation from the chromosomal mass during mitosis (mitotic sorting, Fig. 1A upper panel 9.5 hours, and bottom panel 4.5 hours, SFig. 2I, SFig. 3A (1^st^ example), B). Third, the sorting of plasmid DNA from chromosomal DNA occurs rapidly; plasmid foci formed either during mitosis or in the nucleus during interphase relocated to the cytoplasm within 1 hour after their appearance (median, SFig. 3C). Fourth, in contrast to micronuclei formed by lagging chromosomes or parts of them, plasmid foci formed during mitosis are predominantly formed before (88 % of mitotically appearing plasmid foci) and not during anaphase, when chromosomal fragments or lagging chromosomes become visible as distinct units (SFig. 3D, E). Furthermore, plasmid foci unlike micronuclei never formed in the region between the separating chromosomes during anaphase (Fig. 1A, mitotic formation 4.5 hours, SFig. 3D, E, (Liu et al., 2018; Orr et al., 2021; Wang et al., 2016)). Therefore, we conclude that the dynamics of LacI decorated plasmid DNA are distinct from the mitotic separation of chromosomal fragments or lagging chromosomes from the chromosomal mass. Finally, most plasmid foci are formed during interphase (58 %) and are thus not mitotic products in contrast to micronuclei. Overall, these data reveal that HeLa cells have three ways to specifically sort plasmid DNA away from the chromosomes and that the cell collects plasmid DNA in the cytoplasm where it persists.

### Cytoplasmic plasmid foci remain separated from chromosomes over extended periods of time

Next, we extended the period of live cell imaging up to 122.5 hours to assess if the separation between chromosomes and plasmid DNA is maintained for long time periods (Fig. 1E, F). About 2/3 of the cytoplasmic plasmid foci were maintained and stayed separated from chromosomes during this period, frequently being propagated over 3 cell divisions. The fluorescence of about 1/3 of the foci however decayed over more than 10 hours, before disappearing (Fig. 1F). Since the fluorescence decay occurred only at some plasmid foci within a whole field of view, it was not due to bleaching, but suggests that the DNA was degraded (SFig. 4). Cytoplasmic plasmid foci remained in the cytoplasm during the imaging period and we never observed entry into the nucleus (Fig. 1E, SFig. 4), in contrast to what is reported for DNA of micronuclei (Crasta et al., 2012; Zhang et al., 2015). Thus, even up to 122.5 hours after transfection, plasmid foci behaved similarly to early periods after lipofection. Thus, the separation between chromosomal DNA and plasmid DNA is persistent over several divisions, consistent with (Wang et al., 2016).

### Cells harboring a single cytoplasmic plasmid focus are dominant under diverse conditions

We next studied whether these observations were time- and plasmid type-dependent and occurred in other cell types. We have previously shown that most pLacO lipofected MDCK cells (non-transformed canine kidney cells) predominantly had one cytoplasmic focus per cell, 24 hours after transfection. Here we assessed how MDCK-LacI and HeLa-LacI cells handled plasmid DNA at different times after electroporation and lipofection (SFig. 5A-C). Two critical results are highlighted here. First, in both cell lines and employing both transfection methods, most cells with plasmid foci (grouped into classes of 1- and various multi-foci cells) had a single plasmid focus, regardless of the time point after transfection (3 hours – 72 hours; SFig. 2D, SFig. 5C). Further, the analysis of the electroporation experiments shows that at 3 hours the sum of all different classes of cells with multiple foci pooled together (61 %) is larger than the fraction of cells with only one focus (39 %). This ratio changed over time. Notably, between 24 hours and 72 hours after transfection, the fraction of multi-foci cells decreased strongly, while that of the 1-focus cells increased (1-focus cells: 50 % at 24 hours; 82 % at 72 hours). As this occurred in the absence of further plasmid uptake - in contrast to lipofection - the data suggests that either multi-foci cells died, or a single cytoplasmic focus is differentiated from other plasmid foci in a cell and selectively maintained.

We next tested if the LacO repeat sequence had any effect on the cytoplasmic localization of transfected plasmids. For this, we lipofected plasmids with and without LacO repeats and with or without coding sequences into HeLa cells. Subsequently, we used FISH to visualize these plasmids (SFig. 5D-F). For all tested plasmids, most focus-containing cells had a single cytoplasmic focus 24 hours after lipofection, similar to our experiments where we visualized pLacO with LacI fluorescence (SFig. 5E, F). These results show that plasmid DNA is preferentially maintained in a single plasmid focus in the cytoplasm of mammalian cells, regardless of the cell line, plasmid type, or transfection method.

## Plasmid DNA localizes to the cytoplasm in an ER-enwrapped compartment

The cytoplasmic plasmid foci are ideal to assess if and which membrane is associated with them. As the nuclear envelope originates from the endoplasmic reticulum (ER) and ER is abundant in the cytoplasm, we first quantitatively characterized the association of plasmid DNA with ER. 24 hours after pLacO lipofection into HeLa-LacI cells expressing either the ER transmembrane reporter Sec61-mCherry (Fig. 2A) or the ER-lumen reporter eGFP-KDEL (Fig. 2B), all cytoplasmic plasmid foci colocalized with both ER reporters. Immunofluorescent detection of the ER-residing LEM-domain protein LEM4 (ANKLE2) confirmed the presence of ER at all cytoplasmic plasmid foci 24 hours after lipofection (Fig. 2C) or electroporation (SFig. 6A). The intensities of the ER-reporters KDEL and LEM4 were similar at the plasmid focus compared to the overall ER in 71 % to 91 % of the cases, (category referred to as "non-enriched", Fig. 2B, C, SFig. 6A). However, the intensity of the ER transmembrane marker Sec61 was frequently higher at the plasmid focus compared to the overall ER (category referred to as "enriched" in 61 % of the cases, Fig. 2A, SFig. 6A). Thus, ER membrane and lumenal ER proteins were always present at the cytoplasmic plasmid focus, suggesting that a double membrane encloses the plasmid DNA, reminiscent of the nuclear envelope.

**Fig. 2.**
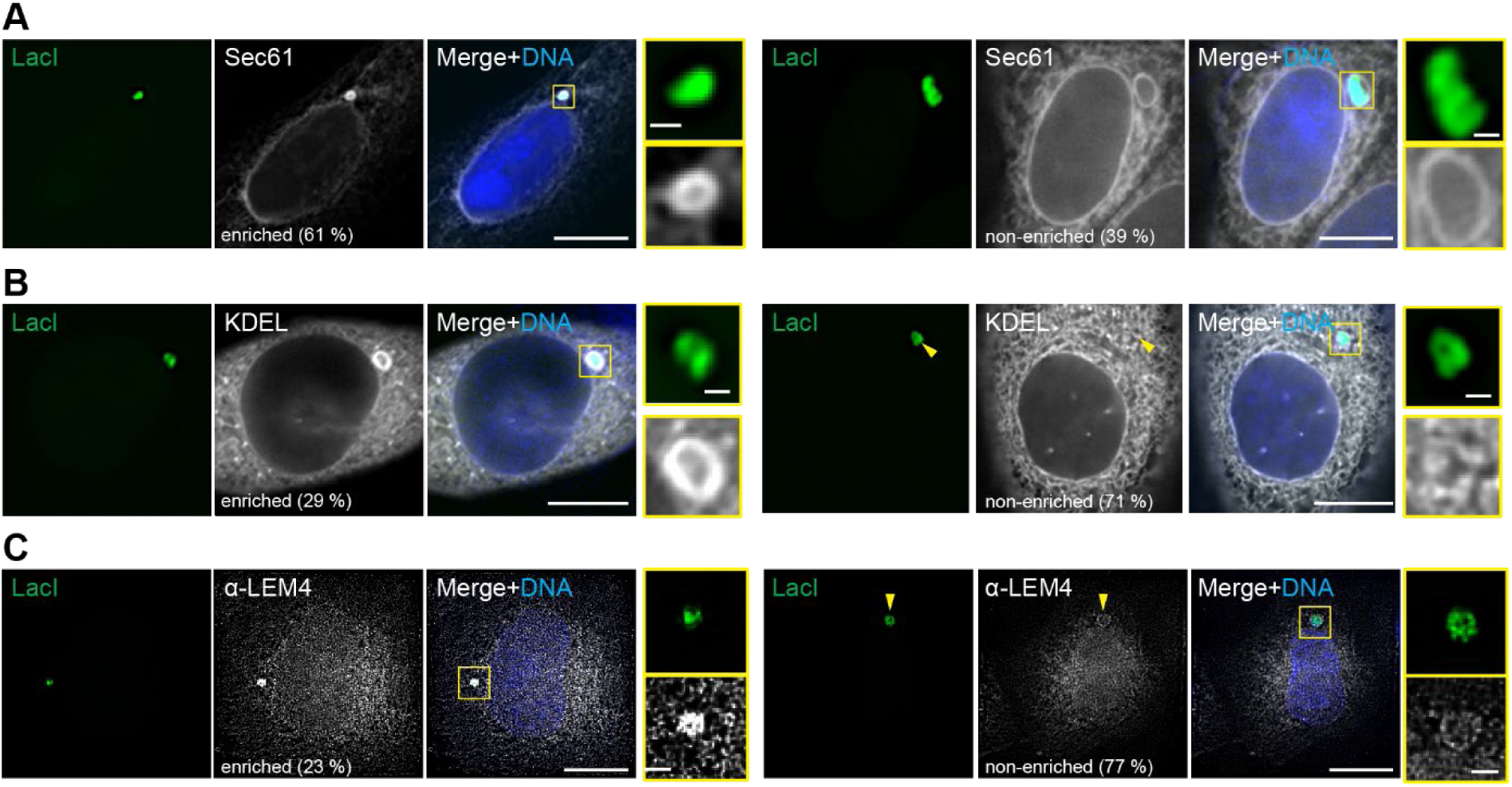
ER enwraps cytoplasmic plasmid DNA. (**A-C**) Representative images of the localization of ER reporters in pLacO transfected HeLa-LacI cells 24 hours after lipofection with the frequency of two localization patterns (enriched and non-enriched, relative to the intensity of the surrounding ER). DNA, blue (Hoechst stain). Single z-slice images, deconvolved. Insets: focus; scale bars: in big images: 10 µm, in insets: 1 µm. (**A**) Transient expression of Sec61-mCherry. Pooled data of 4 exp., total n(cells): 76. n(foci): 79. % relative to all foci analyzed. (**B**) Transient expression of GFP-KDEL. Arrowhead, position of focus; 3 exp.; total n(cells): 80; n(foci): 96. % relative to all foci analyzed. (**C**) Anti-LEM4 immunostaining 24 hours after pLacO lipofection. 2 exp. n(cells): 84; n(foci): 84. % relative to all foci analyzed.

### A special double membrane enwraps cytoplasmic plasmid DNA

To visualize the cytoplasmic plasmid focus at higher resolution, we used correlative light and electron microscopy (CLEM) in interphase HeLa cells 24 hours after pLacO lipofection. In these images, a double membrane enclosing the cytoplasmic plasmid focus is clearly visible (Fig. 3A, yellow arrowheads in the blue inset) similarly to the nuclear envelope (yellow arrowheads in the green inset). The membrane surrounding the plasmid focus has fenestrations, indicating that it may be an open compartment (green arrowheads in the blue inset). Moreover, this membrane connects to the ER (red arrowhead in the blue inset), consistent with the presence of ER proteins at plasmid foci. The cytoplasmic plasmid focus has a higher electron density than the interphase chromosomes in the nucleus suggesting a denser DNA packing in the plasmid focus. Overall, aside from the fenestrations, this compartment’s membrane organization is highly reminiscent of the nuclear envelope surrounding chromosomal DNA.

**Fig. 3.**
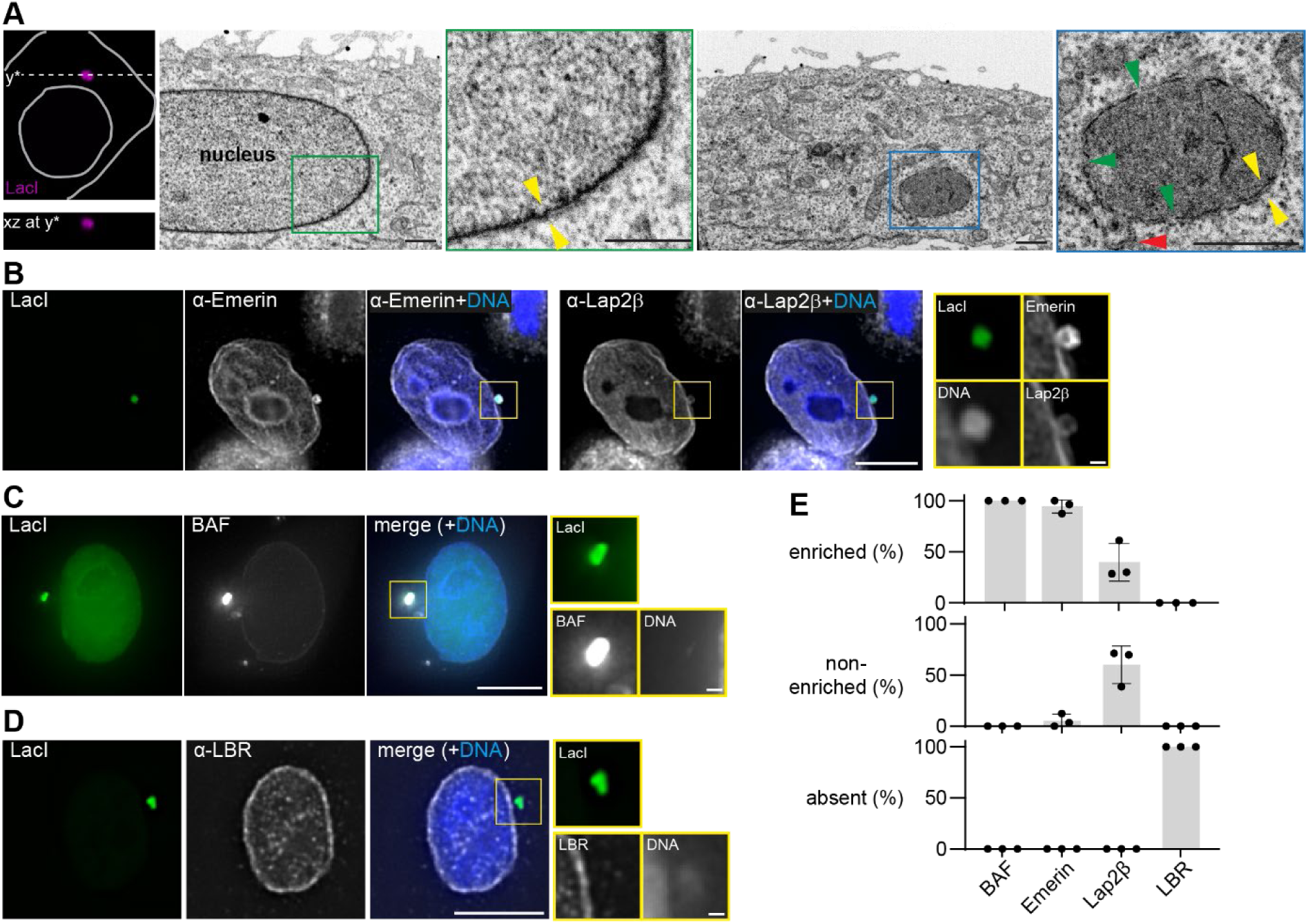
A special double membrane enwraps cytoplasmic plasmid DNA. (**A**) Correlative fluorescence with electron microscopy (CLEM) of an interphase cell containing one pLacO focus. Left images: confocal images; upper: single z-slice xy-image superimposed with a grey cell outline and the y*-cut line; lower: xz-view of the upper cell along the dashed y* line. The first overview EM image depicts a part of the nucleus of the cell shown in the confocal images. The second overview EM image corresponds to the same cell imaged at y*. Insets: focus, blue square; part of the interphase nucleus, green square; double-layered NE, yellow arrowhead pair; membrane connecting proximal ER and the focus, red arrowhead; gaps in the focus envelope, green arrowheads; scale bars 1 µm. (**B-D**) Representative images of the localization of indicated reporters and foci in HeLa-LacI. Images: single z-slice, deconvolved; insets: foci; scale bars: in big images 10 µm; in insets: 1 µm; DNA, blue (Hoechst stain). (**E**) Quantification of relative localization patterns (absent, non-enriched, enriched relative to the NE) of indicated reporters 24 hours after lipofection of pLacO. 3 exp. (circles); mean and SD, each with total numbers: n(Lap2β, foci): 52; n(Emerin, foci): 62; n(LBR, foci): 63; n(LBR, cells): 54; n(BAF, foci): 23; n(BAF, cells): 23.

To further investigate the similarities between the membrane enclosing the plasmid focus and the nuclear envelope, we probed for the presence of inner nuclear membrane proteins at cytoplasmic plasmid foci 24 hours after transfection with pLacO. We paid particular attention to those that could participate in a DNA-membrane tethering at cytoplasmic plasmid foci. One tether at the nuclear envelope is composed of BAF and INM-membrane proteins with a LEM-domain, like Emerin or Lap2β (Ibrahim et al., 2011; Lee et al., 2001; Zheng et al., 2000). Remarkably, cytoplasmic plasmid foci always contained BAF alongside Emerin and Lap2β (Fig. 3B, C, E) (Ibrahim et al., 2011; Kobayashi et al., 2015). BAF was in addition always enriched at plasmid foci compared to the nuclear envelope suggesting a high density of plasmid molecules. More remarkable is that Emerin was enriched at nearly all foci (90 %, Fig. 3E, SFig. 6B). Lap2β was less frequently enriched (40 %; Fig. 3E, SFig. 6B). These results suggest that LEM-domain proteins and BAF might tether plasmid DNA to surrounding ER membranes. Another known tether at the nuclear envelope involves the transmembrane protein Lamin B Receptor (LBR), which binds to heterochromatin protein 1, repressing transcription in the nearby chromatin (Ye and Worman, 1996). LBR was not detected at plasmid foci (Fig. 3D, E). Therefore, we conclude that this second DNA-membrane tether is missing. Overall, the double membrane around the cytoplasmic plasmid focus has both similarities (presence of Emerin and Lap2β) and differences (fenestrations, absence of LBR, enrichment of Emerin) to the nuclear envelope.

### The special double membrane enwrapping cytoplasmic plasmids is devoid of functional NPCs

Next, we probed for NPCs in the membrane around cytoplasmic plasmid foci in HeLa cells, 24 hours after transfection with pLacO. NPCs appear to be absent, as both FG-repeat containing nuclear pore proteins (NUPs; anti-FG repeat) and Embryonic Large Molecule Derived From Yolk Sac (ELYS), which is required for NPC assembly (Rasala et al., 2006), were both absent from cytoplasmic plasmid foci in 94 % of the cases (Fig. 4A). The rare cases when these proteins are present at the plasmid focus might reflect remnants of nuclear budding events. The transmembrane protein Nuclear Envelope Pore Membrane 121 (POM121) was always absent at plasmid foci (Fig. 4B). We also probed for evidence of NPC-mediated nuclear-cytoplasmic transport at cytoplasmic plasmid foci. Here, we observed that the Importin β-binding Domain (IBB-GFP) was always absent (Fig. 4C). In addition, the nucleotide exchange factor for Ran (Regulator of Chromatin Condensation 1, RCC1), which supports NPC formation and establishes a Ran-gradient across the enclosing membrane, was never detected at cytoplasmic plasmid foci (Fig. 4D, (Walther et al., 2003)). We conclude that the double membrane enclosing cytoplasmic plasmid DNA is devoid of functional NPCs.

**Fig. 4.**
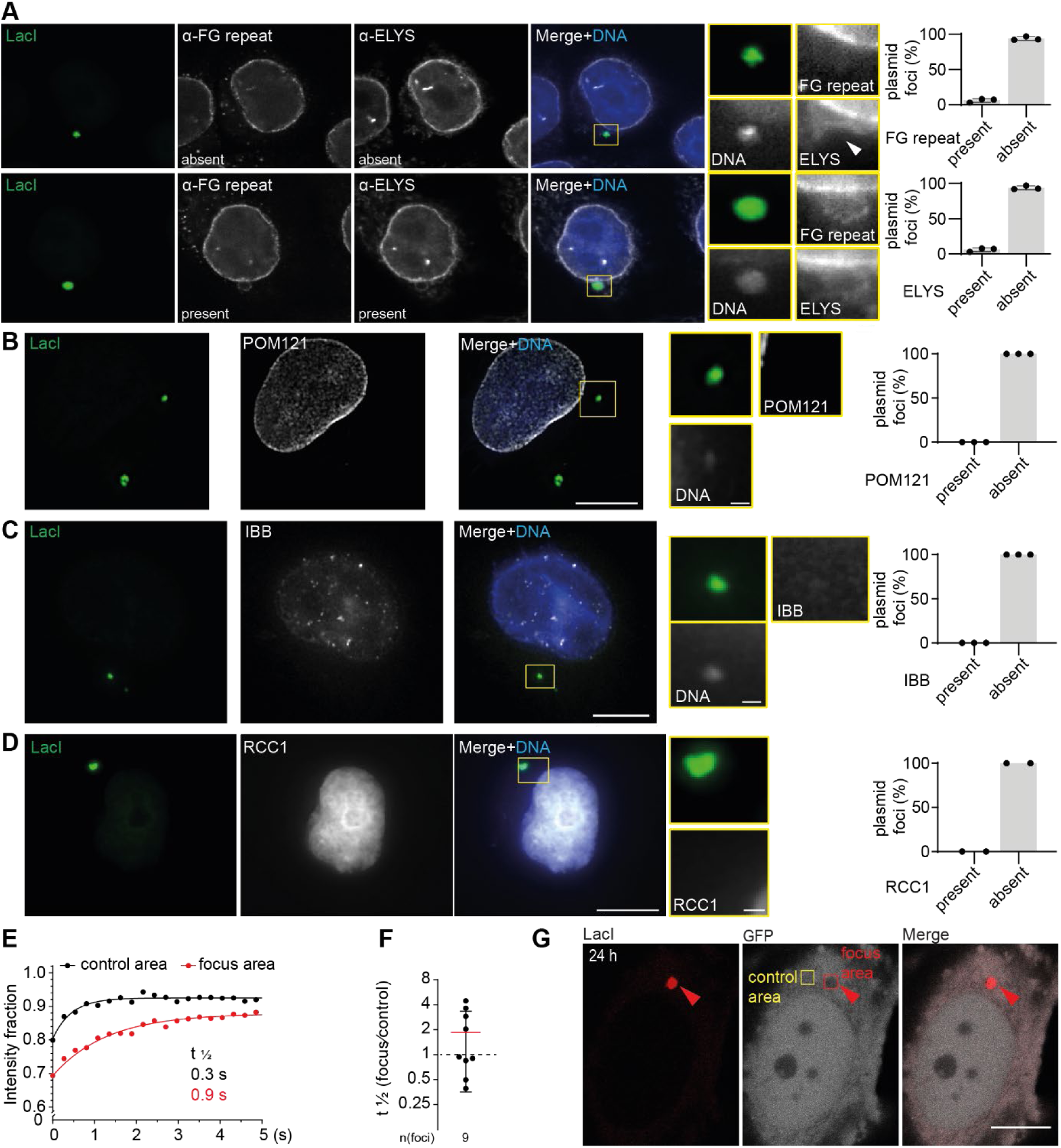
The plasmid enwrapping envelope is devoid of functional NPCs but not closed. (**A-D**) Representative images of HeLa-LacI cells 24 hours after lipofection with pLacO. Insets: focus; scale bars: in big images: 10 µm, in insets: 1 µm; focus, arrowheads. DNA, blue (Hoechst stain). Right quantification; % relative to all foci; 2-3 exp., 1 exp., circle; mean & SD. (**A**) Immunostaining for ELYS and FG-repeats. Upper: absence, lower: presence example, 3 exp., n(ELYS, FG-repeats, foci): 111. (**B-D**) Images single z-slice, deconvolved. (**B**) POM121. 3 exp.; n(foci): 55. (**C**) IBB. 3 exp.; n(foci): 22; n(cells): 17. (**D**) RCC1. 2 exp.; n(foci): 55; n(RCC1, cells): 48. (**E-G**) FRAP analysis in HeLa cells transiently expressing LacI-mCherry and soluble GFP 24 hours after pLacO transfection. (**E**) Recovery of bleached GFP over time; t_1/2:_ recovery time for half of GFP intensity. (**F**) Quantification of (E): Ratio of t_1/2_ at focus area versus control area. Mean & SD; 3 exp.; 1 measurement, circle. n(foci): 9. (**G**) Representative images of bleaching areas. Focus area, red square; control area, yellow area. Scale bar: 10 µm.

The EM analysis indicated that the cytoplasmic plasmid compartment is not entirely closed (Fig. 3A). Therefore, we tested whether soluble GFP could access the cytoplasmic plasmid compartment by Fluorescence Recovery After Photobleaching (FRAP) in HeLa-LacI cells 24 hours after lipofection of pLacO. The GFP fluorescence was bleached at the cytoplasmic plasmid focus and a reference area in the cytoplasm (Fig.4 E-G). Fluorescence recovery took place in both areas, with the recovery time (the time when half of the bleached signal is recovered, t_1/2_) being slightly longer for areas containing the plasmid focus (1.8 times), showing that the diffusion of GFP molecules into the plasmid focus is hindered, but not abolished. Thus, the special double membrane enclosing cytoplasmic plasmid DNA, while devoid of functional NPCs, still allows exchange with the cytoplasm.

### A plasmid focus formed in the cytoplasm is rapidly enwrapped by membrane

The nuclear membrane rapidly encloses chromosomal DNA at the end of mitosis. Therefore, we assayed the time scale of membrane association with cytoplasmic plasmid DNA. We used HeLa cells stably expressing Lap2β-GFP and transiently expressing LacI-mCherry without NLS to ensure enough LacI was present in the cytoplasm, allowing early cytoplasmic plasmid visualization. In addition, the cells were thymidine-synchronized to ensure a higher homogeneity of the cell population. These cells were lipofected with pLacO and imaged every 15 min for 25.25 hours starting after the addition of the lipofection-DNA mix to analyze appearing plasmid foci (Fig. 5). 51 % of such plasmid foci were already associated with membrane at their appearance, as reported by Lap2β (Fig. 5D). Over time, plasmid foci were increasingly associated with Lap2β-containing membrane. Finally, 75 min after appearance, 97 % of the plasmid foci were Lap2β-membrane-associated (Fig. 5D). Thus, membrane association appears to accompany plasmid focus appearance (Fig. 5C, D).

**Fig. 5.**
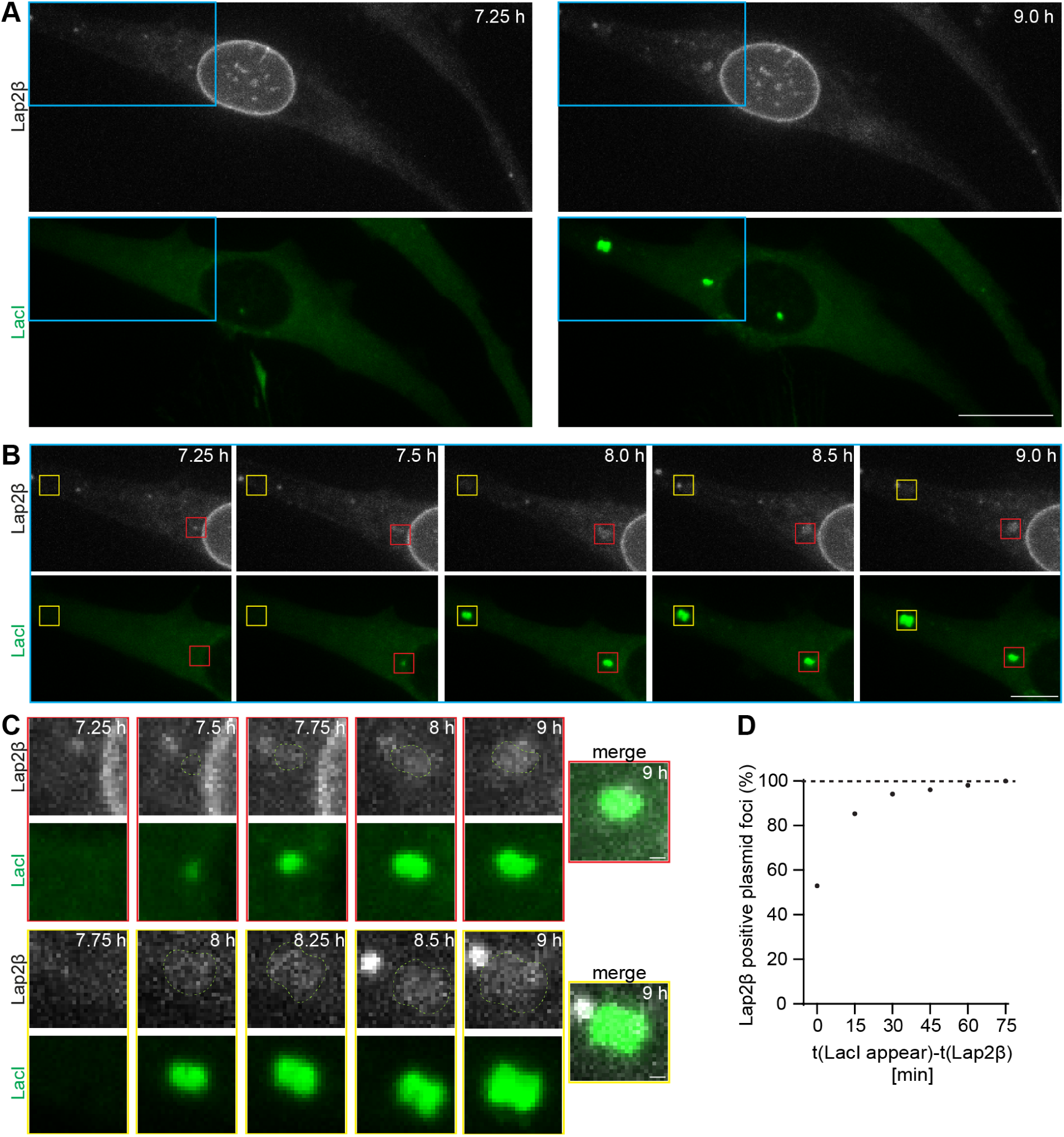
A plasmid focus formed in the cytoplasm is rapidly enwrapped by membrane. (**A-C**) Time-lapse images of HeLa cells stably expressing Lap2β-GFP (Lap2β) and transiently expressing LacI-mCherry (LacI) after lipofection with pLacO. Time, after pLacO transfection. (**A**) Overview images of the cell at 7.25 hours and 9 hours after transfection. The area in the blue square is enlarged in (B). Scale bar, 20 µm. (**B**) Enlarged part of the cell in (A). Focus forms with concomitant Lap2β association, red square. Focus forms with no observable Lap2β association during imaging, yellow square; scale bar, 10 µm. (**C**) Enlarged squares of (B). Focus outline, superimposed dashed line; lower row: Lap2β channel boosted, non-boosted images in (A,B); scale bar, 1 µm. (**D**) Cumulative fraction of foci associated with Lap2β in dependence on the duration of Lap2β association after focus appearance. 1 exp., n(foci): 105; n(cells): 49.

### Emerin is enriched at cytoplasmic plasmid compartments in primary human cells

To probe if the special double membrane can also form around cytoplasmic plasmid foci in non-immortalized cells, we transfected primary human fibroblasts with pLacO and visualized the plasmid with transiently expressed LacI-NLS-GFP. In addition, we immunostained the cells for Emerin or LEM4 (Fig. 6A, B). We noticed that these primary cells divided significantly less frequently than HeLa cells and therefore analyzed the cells 48 hours after pLacO transfection. Also here, most transfected cells had a single plasmid focus (SFig. 6C). Emerin was present at each plasmid focus amongst cells with one plasmid focus and even enriched in 97 % of the instances compared to the surrounding ER or nuclear envelope (Fig. 6A). Also, all plasmid foci in multi-foci cells were Emerin positive apart from one cell, where three foci didn’t colocalize with Emerin, possibly because the reaction to transfection is delayed in these primary cells compared to HeLa cells (SFig. 6D). Remarkably, in multi-foci cells, several plasmid foci of a single cell were Emerin enriched (SFig. 6D, F). Similarly, LEM4 was always present in cells with one plasmid focus and, in 45 % of these cells, even enriched compared to the surrounding ER. This is qualitatively similar to HeLa cells (Fig. 6B). In multi-foci cells LEM4 was always present, except for 2 plasmid foci in two different cells (SFig. 6E).

**Fig. 6.**
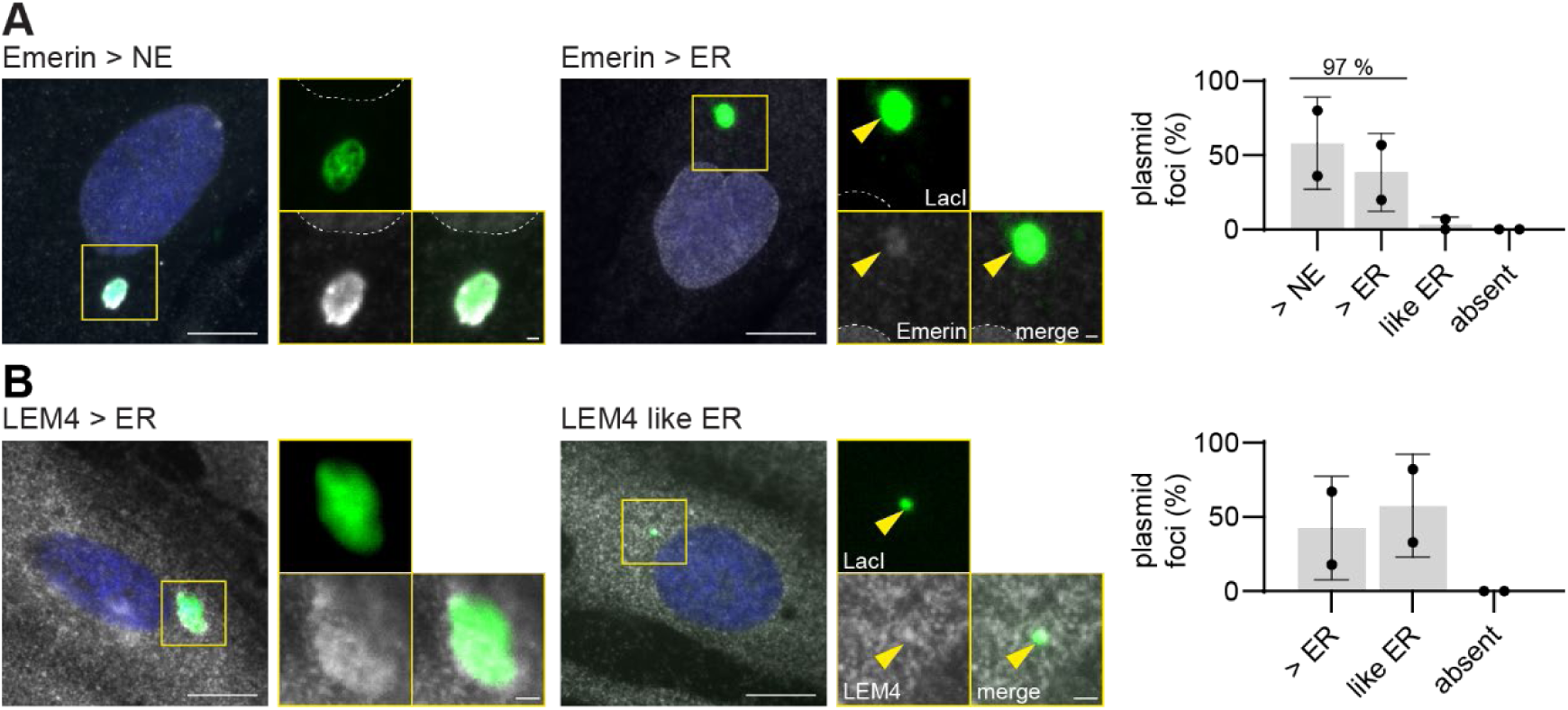
Exclusomes containing plasmid DNA exist in primary human fibroblasts. (**A, B**) Primary human fibroblasts 48 hours after lipofection with pLacO and plasmid encoding LacI-NLS-GFP, immunostained for Emerin (A) and LEM4 (B). Single z-slice images. Insets: focus; scale bars: in big images 10 µm; in insets: 1 µm; DNA, blue (Hoechst stain). (**A**) Representative images of indicated classification. Outline of nucleus, dashed line. Right graph: Emerin’s intensity relative to the NE (> NE) and the ER (> ER, like ER) in cells with 1 cytoplasmic focus. 2 exp., 1 exp., circle; mean and SD; n(foci): 29. (**B**) Representative images for indicated classification of LEM4. Right graph: LEM4 intensity at focus relative to the ER (> ER, like ER) in cells with 1 cytoplasmic focus. 2 exp., 1 exp., circle; mean and SD; n(foci): 38.

Thus, both in primary human fibroblasts as well as in HeLa cells, plasmid DNA is excluded from the nucleus and localizes to the cytoplasm where it exists predominantly in one membranous organelle. We term this organelle the exclusome. A diagnostic hallmark of the exclusome is that Emerin is enriched as compared to the nuclear envelope. The exclusome envelope is further characterized by the presence of fenestrations, presence of Lap2β, presence and even occasional concentration of ER-membrane proteins like Sec61 or LEM4, and the absence of NPCs and LBR.

### Interference with Emerin’s function reduces the compartmentalization of plasmid DNA in the cytoplasm

Since Emerin is enriched at ∼90 % of plasmid foci and the formation of a double membrane is concomitant with plasmid focus formation (Fig. 3B, E, Fig. 5), we speculated that Emerin might tether plasmid DNA to the surrounding membrane to establish an exclusome. As Emerin’s LEM-domain binds BAF and BAF binds DNA (Lee et al., 2001), we aimed to interfere with this molecular linkage. To do so, we chose to interfere with the function of Emerin’s LEM-domain by setting up a competition approach with an excess of the soluble LEM-domain of Emerin. Specifically, we overexpressed either the LEM-domain of Emerin fused to GFP and a nuclear export signal ("GFP-LEM") or soluble GFP ("GFP") (Fig. 7A, left side) as a control in synchronized HeLa-LacI cells. Subsequently, pLacO was electroporated and cells that expressed GFP-LEM or GFP were analyzed (Fig. 7A, right side). The competition was successful for two reasons. First, Emerin was less enriched at the nuclear envelope and more present in the ER in cells expressing GFP-LEM compared to control cells expressing GFP. This suggests that the overexpression of GFP-LEM competed with endogenous Emerin for DNA tethering at the nuclear envelope, thus leading to reduced Emerin retention at the nuclear envelope and re-localization to the ER (SFig. 7B, C). Second, Emerin associated also less frequently with cytoplasmic plasmid foci at two time points after pLacO transfection in cells expressing GFP-LEM compared to the control (Fig. 7B, SFig. 7D).

**Fig. 7.**
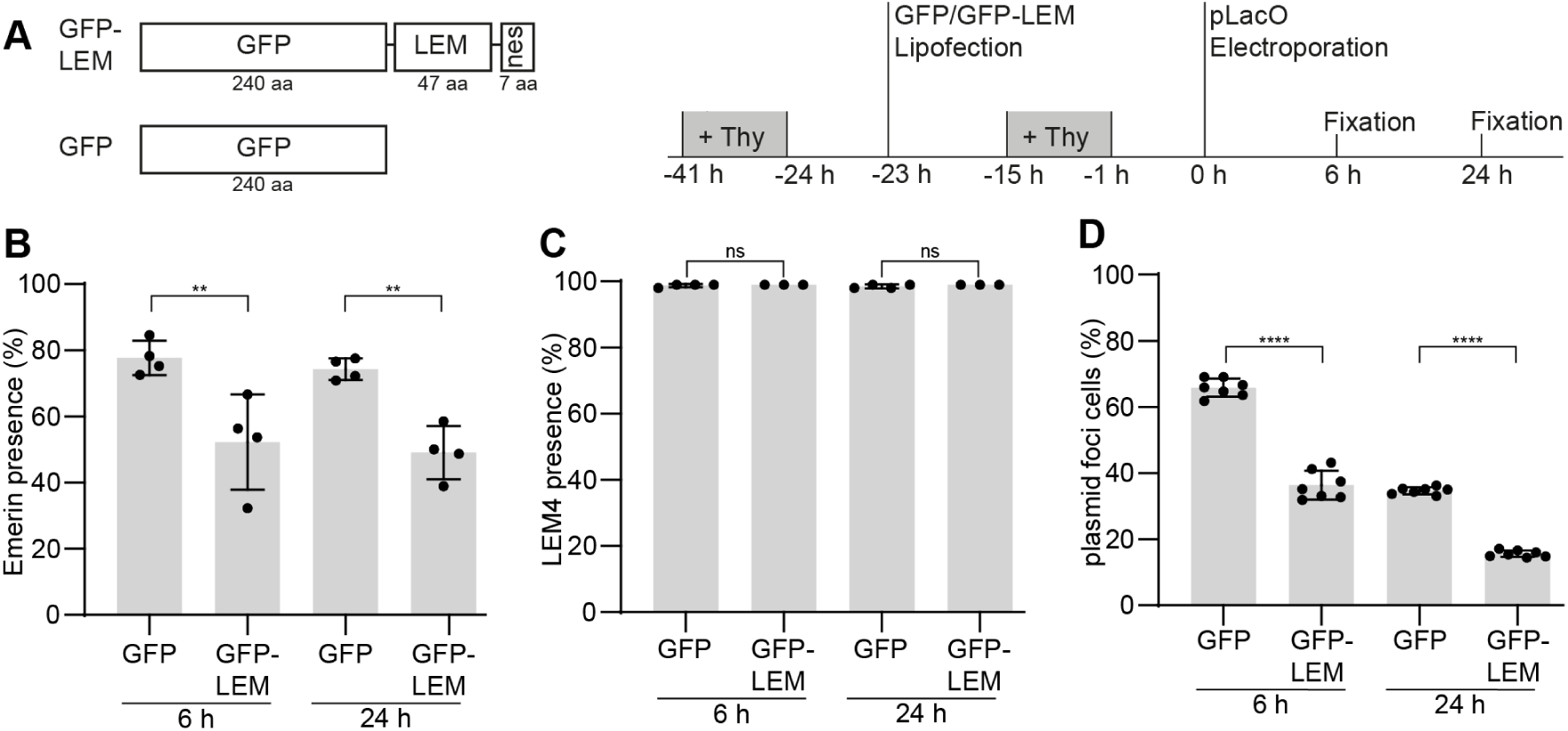
Overexpression of Emerin’s LEM domain reduces cells with plasmid foci. (**A**) Scheme of fusion proteins transiently overexpressed in HeLa-LacI cells (left side) and experimental procedure (right side). GFP-LEM-nes (“GFP-LEM”), soluble GFP (“GFP”); aa: amino acid residues. + Thy: thymidine treatment; time, relative to pLacO transfection. (**B**) Presence of Emerin at cytoplasmic foci 6 hours and 24 hours after electroporation of pLacO. 4 exp., 1 exp., circle; mean and SD; two-way Anova, **: p <0.01; n(foci): GFP 6 hours: 589; GFP 24 hours: 458; GFP-LEM 6 hours: 474; GFP-LEM 24 hours: 413. (**C**) Presence of LEM4 at cytoplasmic foci 6 hours and 24 hours after electroporation of pLacO. 3 to 5 exp. 1 exp., circle; mean and SD; two-way Anova; n.s.: not significant; n(foci): GFP 6 hours: 727; GFP 24 hours: 536; GFP-LEM 6 hours: 334; GFP-LEM 24 hours: 280. (**D**) Frequency of cells containing at least one cytoplasmic focus in GFP and GFP-LEM expressing cells 6 hours and 24 hours after electroporation. 7 exp., 1 exp., circle; mean and SD; two-way Anova; **** p<0.0001; n(cells): GFP 6 hours: 1428; GFP 24 hours: 1460; GFP-LEM 6 hours: 1471; GFP-LEM 24 hours: 1469.

The amino acid sequences of the LEM-domains of Emerin and LEM4 are 44 % similar (SFig. 7A). Because of this, the overexpressed LEM-domain of Emerin might also interfere with the function of other LEM-domain proteins. To test for this possibility and to further characterize the plasmid focus membrane, we probed for the presence of LEM4. The association of LEM4 was not affected in GFP-LEM expressing cells at either 6 hours or 24 hours after pLacO electroporation (Fig. 7C, SFig. 7E). Moreover, LEM4 was present in all conditions at almost all cytoplasmic plasmid foci (99 %, mean of 3 to 5 exp., all conditions), revealing that even Emerin-negative plasmid foci are membrane-enclosed. Furthermore, the presence of LEM4 in Emerin-negative plasmid foci indicates that other proteins might compensate for Emerin’s function, although clearly less efficiently.

Since we could interfere with Emerin’s function, we went on to characterize the effects of GFP-LEM overexpression on the cell’s reaction towards transfected plasmid DNA. We determined how many cells expressing GFP-LEM or GFP, had at least one cytoplasmic plasmid focus. In the GFP-LEM condition, fewer cells contained at least one cytoplasmic plasmid focus compared to control, both 6 hours and 24 hours after pLacO transfection (36 % for GFP-LEM and 66 % for GFP; Fig. 7D). At 24 hours after transfection, the number of cells with plasmid foci were halved in both conditions compared to the 6-hour time point due to cell division and asymmetric partitioning of the plasmid foci (16 % for GFP-LEM and 35 % for GFP). Together, these data suggest that Emerin, through its LEM-domain, supports the compartmentalization of plasmid DNA within the cytoplasm of mammalian cells.

### Exclusomes can contain telomeric DNA

In cells undergoing alternative lengthening of telomeres (ALT), like the osteosarcoma cell line U2OS, circular extra-chromosomal DNA of telomeric origin is abundant (Cesare and Griffith, 2004). Remarkably, in U2OS and several other cancer cell lines, such as WI38-VA13, SaOs2, and KMST-6, between 1 and 4 FISH signals of extra-chromosomal telomeric DNA were detected in the cytoplasm (Chen et al., 2017; Tokutake et al., 1998). In addition, several groups have detected circular extra-chromosomal telomeric DNA in non-ALT cancer cells like HeLa (Regev et al., 1999; Tokutake et al., 1998; Vidaček et al., 2010; Wang et al., 2004). Therefore, we decided to test whether this extra-chromosomal DNA of endogenous origin is membrane-enclosed in the cytoplasm and whether this membrane shares similarities with the nuclear envelope or the exclusome.

We performed FISH experiments in U2OS and HeLa cells using two different fluorescently tagged telomeric probes (TelC and TelG). In all cases, we observed numerous FISH signals in the nuclei and few in the cytoplasm (Fig. 8A, B, SFig. 8A, B). We also demonstrated that the cytoplasmic Tel FISH signals were not artifacts caused by clustered probes as cells simultaneously hybridized with both a telomeric probe as well as a scrambled probe only showed telomeric probes signals (SFig. 8A-C). Further, both types of cytoplasmic telomeric FISH signals indeed labeled DNA, as a DNaseI treatment prior to probe hybridization abolished the signal (SFig. 8D, E).

**Fig. 8.**
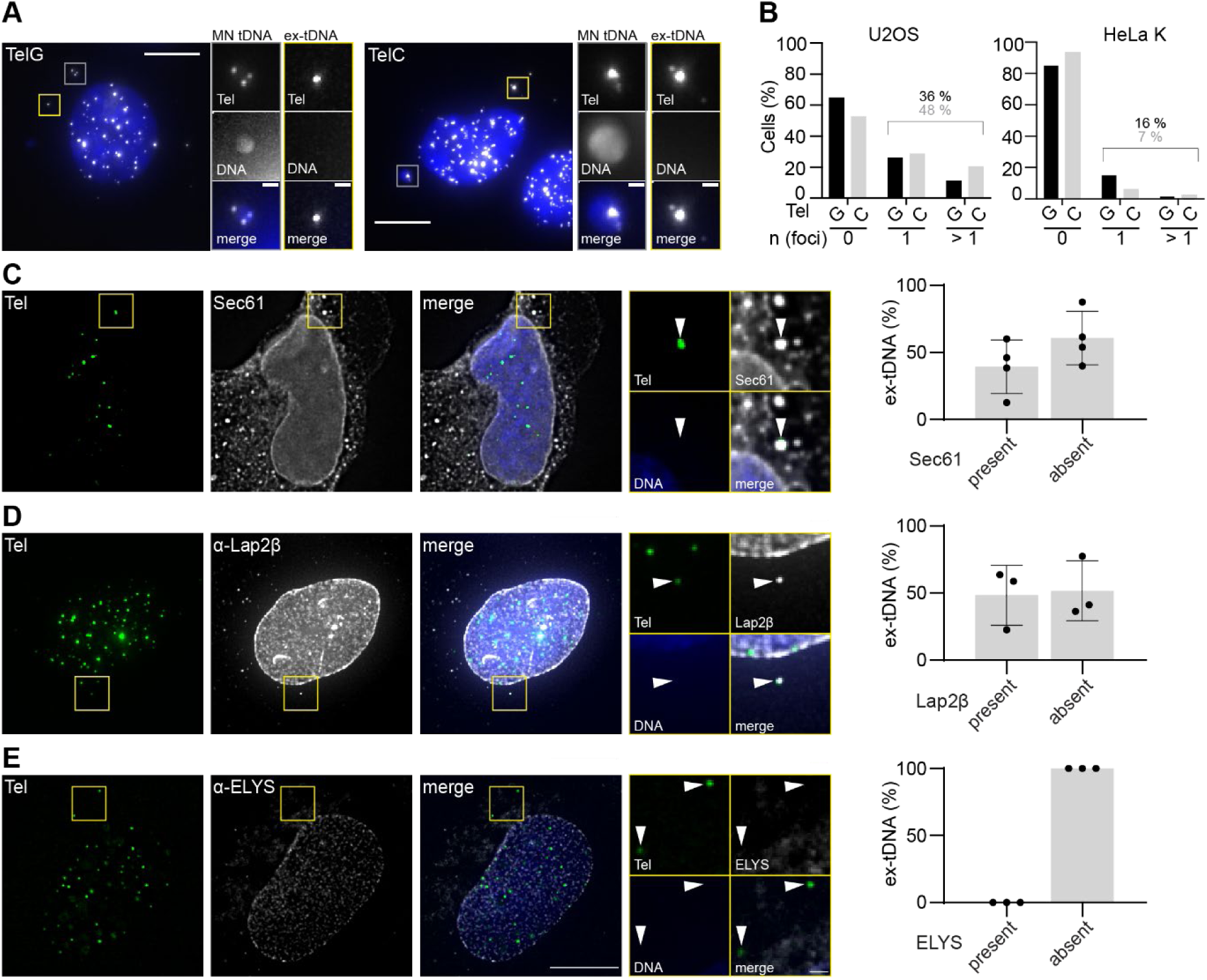
A special envelope enwraps also cytoplasmic extra-chromosomal telomeric DNA. (**A**) Representative images of U2OS cells FISH stained with TelG and TelC probes. Images max. projected, insets: area with ex-tDNA, yellow squares; area with MN tDNA, gray squares. Scale bars: in big images: 10 µm; in insets: 1 µm. DNA, blue (Hoechst stain). (**B**) Frequency of HeLa K and U2OS cells with none, one (1), or more than one (>1) ex-tDNA focus relative to the total cells analyzed. Pooled data of FISH experiments. U2OS-TelG probe: 3 exp.; > 47 cells per exp., total n(cells): 217; U2OS-TelC probe: 3 exp.; > 49 cells per exp., total n(cells): 171; HeLa-TelG probe: 3 exp.; > 55 cells per exp., total n(cells): 210; HeLa-TelC probe: 3 exp.; > 63 cells per exp., total n(cells): 209. (**C-E**) Representative single z-slice images of FISH-IF stained U2OS cells depicting the localization of the reporter proteins (left); quantification of colocalization of respective marker at ex-tDNA focus (right, plot). Big images: max. projected deconvolved; arrowheads, ex-tDNA foci; Areas of tDNA foci, insets. Scale bars: big images: 10 µm, insets: 1 µm; 3 exp.; DNA, blue (Hoechst stain). Signals of TelG probe, overexpressed of Sec61-mCherry (C) and indicated antibodies (D,E). % relative to all ex-tDNA foci analyzed. Mean & SD. Sec61, 4 exp., 1 exp., circle, n(Sec61, ex-tDNA): 95; Lap2β, 3 exp. 1 exp., circle, n(Lap2β, ex-tDNA):1 54; ELYS, 3 exp. 1 exp., circle, n(ELYS, ex-tDNA): 66.

Micronuclei with chromosomal fragments frequently occur in cancer cells. To exclude such compartments from our analysis, we chose Hoechst fluorescence as a criterium, as the median size of circular ex-tDNA is only 5 kb (Cesare and Griffith, 2004). Based on this criterium, we distinguished two types of tDNA in the cytoplasm of both HeLa and U2OS cells: tDNA in Hoechst positive foci, termed <u>m</u>icro<u>n</u>uclear tDNA (MN tDNA, Fig. 8A (grey squares)) as they might contain chromosomal DNA fragments with telomeres; and extrachromosomal tDNA without Hoechst stain, termed ex-tDNA. In the following analyses, we focused on the cytoplasmic ex-tDNA (Fig. 8A (yellow squares)). Overall, there were fewer cells with ex-tDNA foci in the HeLa population (7 - 16 %) compared to the U2OS population (36 - 48 %, SFig. 8C). Such a difference is expected for non-ALT cells versus ALT-cells and is consistent with the notion that ex-tDNA foci represent circular ex-tDNAs. Remarkably, both HeLa and U2OS cells mostly contained one ex-tDNA focus per cell for both types of probes (U2OS 25 %, HeLa 5 %), which is strikingly similar to our findings with transfected plasmid DNA ((Wang et al., 2016), Fig. 8B).

Next, we analyzed whether cytoplasmic ex-tDNA foci are membrane-enclosed in U2OS cells. We could not directly probe for the presence of Emerin, as none of our anti-Emerin antibodies sustained the conditions used in FISH-IF experiments. However, overexpressed Sec61-mCherry colocalized with 47 % of ex-tDNA foci but always colocalized with MN tDNA (Fig. 8C, SFig. 8F). Also, Lap2β was present at 41 % of ex-tDNA foci but always present at MN tDNA foci (Fig. 8D, SFig. 8F). ELYS was never present at ex-tDNA foci but was typically present at MN tDNA (2 out of 3 cases) (Fig. 8E, SFig. 8F). Thus, the co-localization frequencies for the tested ER- and INM-proteins at ex-tDNAs were lower as for plasmid foci, possibly due to the reduced focus size (Fig. 8A, C-E, yellow squares) and the harsh conditions applied during FISH. Notably, NPCs were absent from ex-tDNA but not from tDNA in micronuclei (Fig. 8E, SFig. 8F), consistent with the possibility that the latter are formed during mitosis and have different contents than ex-tDNA foci. Collectively, these results indicate that ex-tDNA can also be contained in exclusomes.

### Both plasmid DNA and ex-tDNA cluster in an exclusome

Due to the observed similarities between ex-tDNA and plasmid DNA, we tested if they colocalize within the same exclusomes. U2OS cells were fixed and immunostained for Lap2β 24 hours after lipofection with pLacO. In addition, the cells were hybridized *in situ* with probes for both LacO and TelC. Indeed, plasmid and ex-tDNA colocalized in the cytoplasm in one Lap2β containing membrane compartment (Fig. 9A). Among all cells with both types of DNA in the cytoplasm (co-existence cells), 26 % had both DNA types in a single cytoplasmic compartment (Fig. 9B, single exp. SFig. 8G). 75 % of such compartments contained Lap2β in their envelope (Fig. 9A, C, SFig. 8H). In 74 % of such Lap2β-containing compartments plasmid and tDNA foci were Hoechst positive, which could represent chromosomal fragments with telomers together with plasmid DNA (Fig. 9C, SFig. 8H). However, the fact that 26 % of such compartments were Hoechst negative reveals that ex-tDNA, and not telomeric DNA from chromosomal ends, colocalized with plasmid DNA in one cytoplasmic membrane-bound compartment (Fig. 9A). Therefore, we conclude that extra-chromosomal DNAs of different origins, such as endogenous telomeric DNA and exogenous plasmid DNA, can cluster in one exclusome (Fig. 9D).

**Fig. 9.**
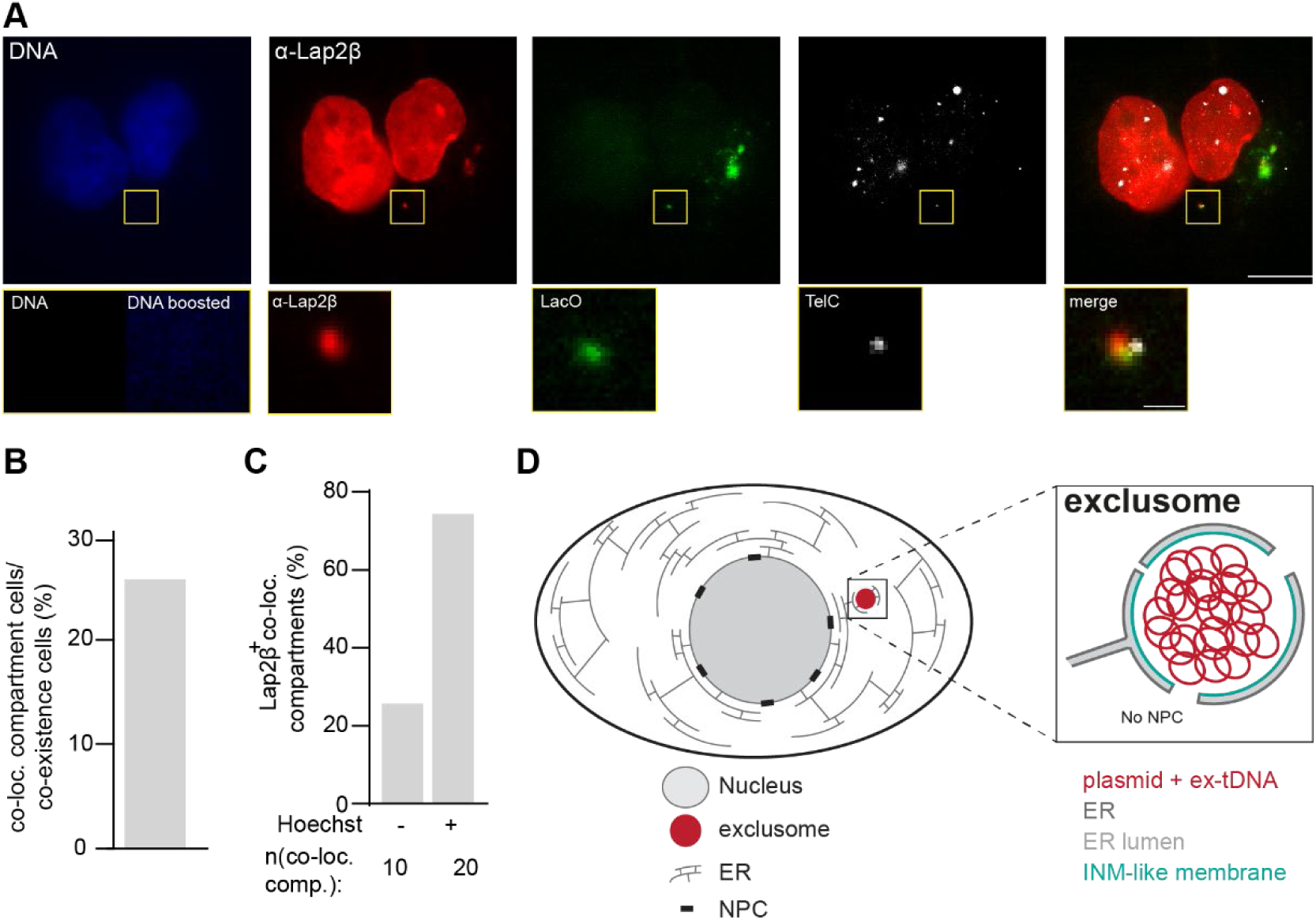
Plasmid DNA and ex-tDNA can be clustered in one exclusome. (**A**) Representative image of a U2OS cell transfected with pLacO, fixed, hybridized with TelC and LacO probes and immunostained against Lap2β. DNA, blue (Hoechst stain). Inset: cytoplasmic compartment with two DNA species; DNA stain inset boosted, DNA boosted. Scale bars: in big images: 10 µm, in insets: 1 µm. (**B**) Frequency of U2OS cells with minimally one co-localization compartment in cells, which contain both DNA species (co-existence cells) 24 hours after pLacO transfection. Pooled data of 3 exp., individual exp. in SFig. 8G. >15 co-localization compartments/exp.; total n(co-existence cells): 155; >3 co-existence cells/exp; total n(co-existence cells): 40. (**C**) Frequency of Lap2β positive cytoplasmic co-localizing compartments. 24 hours after pLacO transfection. Pooled data of 3 exp.; individual exp. in SFig. 8H. >3 co-localizing compartment/exp., total: n(Lap2β+ co-localizing compartment): 30. (**D**) Model of an exclusome in an interphase cell. Overview of a cell with nucleus, ER and exclusome (inlet: the magnified exclusome with details).

## Discussion

Our study reveals that somatic vertebrate cells maintain transfected DNA in a new structure, which we term exclusome and which represents a third cytoplasmic DNA compartment besides the nucleus and mitochondria. The exclusome, mostly one per cell, is a cytoplasmic membranous compartment into which the cell sorts and where it retains transfected plasmid and extra-chromosomal elements, such as telomeric DNA, likely in circular form, for extended periods of time. 24 hours after transfection, the envelope of the exclusome strikingly resembles the nuclear envelope in some aspects but differs from it in others (Fig. 9D). Thus, a nuclear-like envelope was assembled around cytoplasmic plasmid DNA. We observed this in the variety of cells studied here (HeLa, U2OS, MDCK, primary human fibroblasts), suggesting it is not cell type specific amongst somatic cells. This is remarkable, as it enlarges our understanding of the formation of the nucleus and the nuclear envelope.

The exclusome and the nucleus have three striking similarities. Both contain DNA and exist generally each as a single unit in the cell. In both cases their membranous envelope is a sheet-like double membrane derived from the ER (as shown by the presence of Sec61, LEM4, and KDEL-GFP) and comprising specific INM proteins (Lap2β and Emerin). These points suggest that bringing DNA together in one unit and enwrapping it with a minimal ER-derived double membrane containing Lap2β and Emerin might represent a default response to DNA (Kobayashi et al., 2015).

However, there are also clear differences distinguishing the exclusome from the nucleus: its envelope structure and its DNA content. The envelope of the exclusome differs from that of the nucleus, as LBR and NPCs are missing, Emerin is enriched, and membrane fenestrations are present in the exclusome envelope but not in the nuclear envelope. At the end of mitosis, Lap2β and Emerin, are known to arrive within the first membrane patches that assemble on the decondensing chromosomes (Haraguchi et al., 2008). In contrast, complete NPCs are only very late in telophase in the membrane wrapping around the chromosomes (Haraguchi et al., 2000; Otsuka et al., 2018; Otsuka et al., 2023). Similarly, the sealing of fenestrations does not occur in the envelope of the exclusome however it does occur during the formation of the nuclear envelope late in mitosis (Ventimiglia et al., 2018). Thus earlier, but not later steps of nuclear envelope formation concur to generate the exclusome. Remarkably plasmid DNA competed in *in vitro* assays with nuclear envelope formation around chromosomes (Ulbert et al., 2006). Therefore, it could become a useful model for identifying the triggers establishing the contacts between protein containing membrane and DNA, the first steps of nuclear envelope assembly.

Another difference is the enrichment of Emerin in the envelope of the exclusome compared to the nuclear envelope (Haraguchi et al., 2022; Ibrahim et al., 2011; Kobayashi et al., 2015). This hallmark characteristic is in striking contrast to micronuclei, which contain chromosomal fragments or entire chromosomes, as their surrounding membrane is enriched for Lap2β but not for Emerin (Liu et al., 2018). Emerin’s enrichment might be due to liberation of Emerin from the INM as seen upon stress (Buchwalter et al., 2019), or is due to accumulation of ER membrane with newly synthesized Emerin at the site of exclusome formation, similarly as seen for the envelopment of chromosomal fragments (Ferrandiz et al., 2022). Our results further suggest that Emerin and its LEM-domain play a key role in the envelopment of plasmid DNA in the cytoplasm. What drives Emerin specificity for exclusomes, in contrast to LEM4 for example, remains to be investigated. Intriguingly the DNA-viruses, vaccinia virus and mimivirus, generate replication factories from the ER in the host cell cytoplasm (Greseth and Traktman, 2022; Mutsafi et al., 2013). It will be interesting to determine whether the ER at these factories shows similarities to that at the exclusome.

What do these differences possibly mean? Besides NPCs, we noticed that RCC1 and ELYS were both absent from exclusomes. Both regulate early steps of NPC formation (Gómez-Saldivar et al., 2016; Walther et al., 2003) and their absence might explain why exclusomes do not assemble NPCs. The absence of ELYS would explain that of LBR from exclusomes as well (Mimura et al., 2016). Thus, it will be interesting to determine what causes the absence of ELYS and RCC1 from exclusomes. Possibly, it is due to differences in chromatinization of the DNA in the exclusome compared to that of chromosomes in the nucleus, to which RCC1 binds (Chen et al., 2007). Remarkably, human artificial chromosomes (HAC) relocalize to a cytoplasmic “nanonucleus” upon inactivation of their engineered centromere (Nakano et al., 2008). These nanonuclei have not been further characterized. Yet, it is tempting to speculate that the absence of a centromere on a DNA molecule might be signal to sort that DNA into an exclusome. Future studies will determine the molecular determinants governing how a given DNA molecule is enveloped and thus how the cell distinguishes extra-chromosomal DNA like plasmid and ex-tDNA from the chromosomes.

The content of the exclusome differs from that of the nucleus. We reveal processes, by which the cell actively separates chromosomal from extra-chromosomal DNA. The cell sorts transfected plasmid DNA and ex-tDNA into the exclusome, whereas it bundles the chromosomes into the nucleus at the end of mitosis. We show that sorting of incoming plasmid DNA likely occurs directly in the cytoplasm, as most plasmid foci are formed in this compartment. Thereby, only little plasmid DNA reached the nucleus under our transfection and visualization conditions. In this regard, the exclusome prevented contact between the transfected DNA and the chromosomal DNA. When the transfected DNA reached the nucleus, it did not remain there, but was expelled. We identified two modalities by which nuclear plasmids became separated from chromosomal DNA, which remove plasmid from the nucleus, where it is generally thought to be expressed. The first one, mitotic sorting, expels plasmid DNA to the cytoplasm during mitosis. This mechanism was also reported for plasmid DNA microinjected into the nucleus (Ludtke et al., 2002). The second one occurs during interphase and involves the budding of a newly formed exclusome out of the nuclear envelope, into the cytoplasm. Similar, chromosome-derived large circular DNAs encoding c-myc visualized by FISH localized in nuclear buds in the human colorectal adenocarcinoma cancer cell line COLO 320DM (Shimizu et al., 1998). In all cases, the transfected DNA and even originally nuclear ex-tDNA end up in a cytoplasmic exclusome. Thus, the cell has a machinery distinguishing and sorting these DNAs from chromosomal DNA.

Cells maintain an exclusome with plasmid DNA for long periods of time over multiple cell divisions, but the physiological relevance of this is unknown. Generally, cytoplasmic DNA is sensed as a danger signal by the cyclic guanosine monophosphate–adenosine monophosphate synthase (cGAS), which provokes type I interferon production to warn neighboring cells about this rogue DNA. The organismal immune system would subsequently remove such a cell. As cGAS was found at transfected cytoplasmic plasmid DNA, the exclusome might be an immunologically relevant signaling hub until the cell is eliminated (Guey et al., 2020). Additionally, an exclusome might alter the cellular reaction to other incoming DNAs and could explain why transfection or virus infection are less efficient subsequently to a first transfection (Grandjean et al., 2011; Langereis et al., 2015). In such a scenario, we suggest that the exclusome might act as a memory deposit for both systemic and cell-autonomous immunity towards DNA. In line with this, DNA enwrapping seems to be in competition with nucleases, as most of the transfected DNA is likely degraded before being captured in an exclusome (Shimizu et al., 2005). Also, plasmid foci occasionally disappeared over time, indicating that the maintenance of an exclusome is constantly challenged by cellular defense processes.

To conclude, we identified that cells distinguish, sort, and cluster extra-chromosomal DNAs away from their chromosomes into a membranous compartment in the cytoplasm, the exclusome. The envelope of the exclusome bears some similarities to the nuclear envelope but also differences as it e.g., does not perform the NPC-controlled nucleo-cytoplasmic exchange of the nuclear envelope. Remarkably, most exclusomes form in interphase cells, whereas the nucleus of mammalian cells forms specifically at mitotic exit. This suggests that DNA clustering and the steps of nuclear envelope formation that are common to the exclusome and the nucleus are not dependent on cell cycle regulation. We suggest that they may have evolved together with open mitosis as a mechanism to exclude extra-chromosomal DNA from the nucleus. Indeed, following transfected plasmid DNA in budding yeast, undergoing closed mitosis, revealed that exclusomes are not present (Denoth-Lippuner et al., 2014; Shcheprova et al., 2008). Still, is the exclusome biology conserved in other, especially non-vertebrate, organisms? We expect it to be especially prominent in organisms undergoing open mitosis and in which cGAS or an analogous system is present.

## Material and methods

### 1. Mammalian cell lines

All cell lines listed in the following were cultured at 37 °C with 5 % CO_2_ in a humidified incubator in the indicated media.

#### 1.1. HeLa

HeLa Kyoto (HeLa K) (internal Lab ID: MMC278), human cervical cancer cells were a kind gift from P. Meraldi (ETHZ, Switzerland) and originated from S. Narumiya, (Kyoto University, Japan). Cells were cultured in Dulbecco’s modified medium (DMEM) with high glucose (Gibco; Thermo Fisher Scientific, Basel, Switzerland) plus 10 % FCS (PAA Laboratories; Pasching, Austria) and P/S (100 U/ml penicillin and 100 mg/ml streptomycin; Gibco, Thermo Fisher Scientific). Results with HeLa K are shown in Fig. 8A, B, SFig. 8A, C.

HeLa K cells stably expressing a mutant form of LacI that cannot tetramerize in the fusion proteins LacI-NLS-mCherry (internal Lab ID: MMC114) and LacI-NLS-EGFP (EGFP, enhanced GFP, internal Lab ID: MMC105) were described in (Wang et al., 2016). Cells were cultured as Hela K cells, but the medium contained in addition 5 µg/ml Blasticidine S (Gibco). Both LacI-NLS-XFP cell lines, as well as HeLa transiently expressing LacI-mCherry are referred to “HeLa-LacI” throughout the result and legend text. Results with HeLa LacI-NLS-EGFP are shown in Fig. 2A, C, Fig. 3B, SFig. 1, SFig. 4, SFig. 5C, E, F. Results with HeLa LacI-NLS-mCherry are shown in Fig. 2B, Fig. 3A, C, Fig. 4A-D, Fig. 7, SFig. 1, SFig. 7. HeLa K cells transiently expressing LacI-mCherry were used in Fig. 4E-G, Fig. 5.

HeLa K cells stably expressing LacI-NLS-EGFP which were in addition MOCK electroporated, is termed ”Control HeLa” (internal Lab ID: MMC248) in SFig. 1 but otherwise “HeLa-LacI” to facilitate readability. Cells were cultured like HeLa K cells, but the medium contained additionally 5 µg/ml Blasticidine S (Gibco). Results with this cell line are shown in Fig. 1, SFig. 1, SFig. 2, SFig 3.

HeLa K cells stably expressing aa 244–453 of Lap2β-GFP (internal Lab ID: MMC84), were kindly provided by U. Kutay and originated from (Mühlhäusser and Kutay, 2007). Results with this cell line are shown in Fig. 5. Expressed aa 244–453 of Lap2β-GFP is termed “Lap2β”.

#### 1.2. MDCK

MDCK II (Madin-Darby canine kidney) cells stably expressing LacI-NLS-EGFP (internal Lab ID: MMC100), hereafter termed MDCK-LacI, were described in (Wang et al., 2016). Results with this cell line are shown in SFig. 5A-C.

#### 1.3. U2OS

U2OS osteosarcoma cells (internal Lab ID: MMC95), were a kind gift from C. Azzalin (Instituto de Medicina Molecular, Portugal, cells originated from A. Londono Vallejo). Cells were cultured in Dulbecco’s modified medium (DMEM) with high glucose (Gibco) plus 10 % FCS (PAA Laboratories), P/S (Gibco). Results with this cell line are shown in Fig. 8, Fig. 9, SFig. 8B-H.

#### 1.4. Primary human fibroblasts

Human primary foreskin fibroblasts (internal Lab ID: MMC281) were kindly provided by Dr. Hans-Dietmar Beer, University of Zurich, Switzerland. The foreskin had been collected with informed written consent of the parents in the context of the Biobank project of the Department of Dermatology, University of Zurich, and its use had been approved by the local and cantonal Research Ethics Committees. Cells were cultured in Dulbecco’s modified medium (DMEM) with high glucose (Gibco) plus 10 % FCS (PAA Laboratories) and P/S (Gibco). Results with these cells used at passage number 6 and 7 are shown in Fig. 6, SFig. 5C-F.

#### 1.5. Cell cycle synchronization

HeLa cells were synchronized using a double thymidine (2 mM, Sigma Aldrich; St. Lewis, Missouri) treatment. Cells were treated with thymidine for 16 hours, released for 8 hours and treated with thymidine a second time for 20 hours. 1 hour after the second thymidine release, pLacO transfection was performed. 6 hours after pLacO transfection cells were washed with 20 U/ml heparin in PBS (3 x 3 min, 37°C). This procedure was used in: Fig. 2A, B, Fig. 3B, D, Fig. 4B-E, Fig. 5.

In a second thymidine treatment protocol, cells were treated with thymidine (2 mM, Sigma Aldrich) for 16 hours, then released for 8 hours. 1 hour after this release, plasmids (GFP, GFP-LEM) were lipofected. Cells were exposed to a second thymidine treatment (2 mM) for 18 hours. Cells were electroporated with pLacO 2 hours after the second thymidine release. This procedure was used in Fig. 7 and SFig. 7.

### 2. Plasmid

#### 2.1. Oligonucleotides used

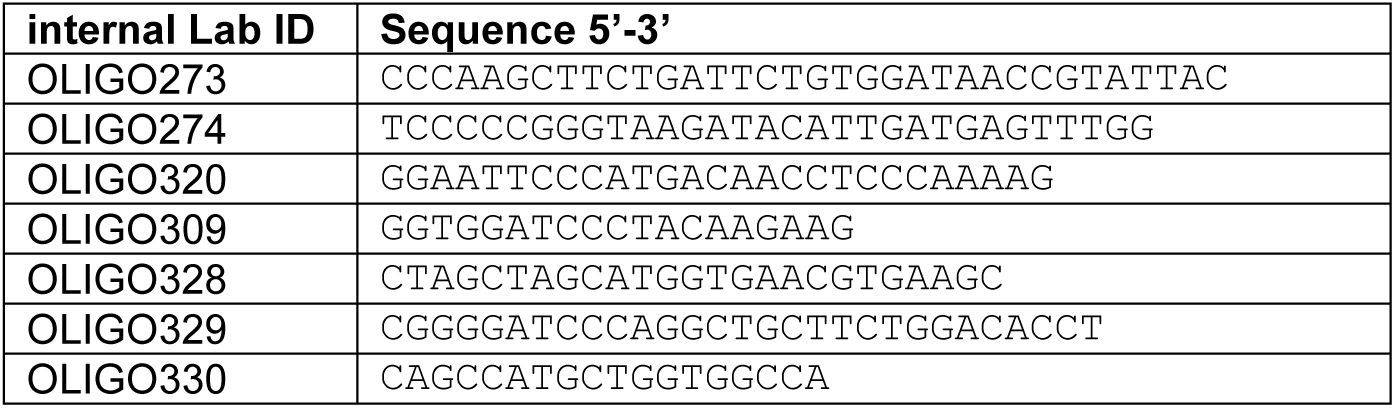

#### 2.2. Plasmid preparation

Plasmid DNA was extracted from *E. coli* bacteria (XL1Blue strain or DH5α strain) using plasmid extraction kits (QIAGEN; Venlo, Netherlands or Macherey Nagel; Düren, Germany). The DNA was purified using either Phenol/Chloroform/Isopropanol, ethanol, or 2-Propanol purification. The purified DNA pellet was resuspended in ddH_2_O of appropriate volume. Plasmid concentration was measured by a NanoDrop Spectrophotometer (Thermo Fisher Scientific).

#### 2.3. Construction of plasmids

pControl 1 is also termed pSR9vector-CMV-mCherry (internal Lab ID: PLA1036): CMV-mCherry-SV40-PA was amplified via PCR and the OLIGO273 and OLIGO274 from p-mCherry-N1 without multiple cloning site (modified Clontech, Takara; United States. Internal Lab ID: PLA1029). The PCR product for CMV-mCherry was cloned into the backbone of pLacO (internal Lab ID: PLA977), after removing the LacO repeats.

pLacI-mCherry (no NLS) (internal Lab ID: PLA1107): LacI was amplified with PCR from LacI-NLS-mEGFP (internal Lab ID: PLA978) using OLIGO328 and OLIGO329 with restriction sites for NheI and BamHI. The backbone vector pIRESpuro2-FLAG-mCherry (internal Lab ID: PLA768, kindly gifted from Yves Barral (IBC, ETH Zurich, Switzerland)) was digested with NheI and BamHI and LacI PCR insert was ligated. Clones were checked with sequencing using OLIGO330.

pEGFP-LEM-nes (internal Lab ID: PLA1098): The sequence of human Emerin’s LEM-domain with a nuclear exclusion signal (nes)(GGAATTCCTCCGAAGATATGGACAACTACGCAGATCTTTCGGATACCGAGCTGACCACCTTGCTGCGCCGGTACAACATCCCGCACGGGCCTGTAGTAGGATCAACTCGTAGGCTTTACGAGAAGAAGATCTTCGAGTACGAGACCCAGAGGCGGCGGGCCCGGGATTTAGCCTTGAAATTAGCAGGTCTTGATATCTACCCCGAAGATTAAGCGGCCGCTAAACTAT) (internal Lab ID: SYN2) was ordered from Lifetechnologies AG (Basel, Switzerland) and inserted into a modified version of pEGFP-N1 (internal lab ID: PLA328).

pEGFP-BAF (internal Lab ID: PLA1089): BAF was amplified by PCR from pEGFP-HIS-BAF (internal Lab ID: PLA1080; was a kind gift from Tokuko Haraguchi (National Institute of Information and Communications Technology 588-2 Iwaoka, Iwaoka-choNishi-ku, Kobe 651-2492, Japan)) with the primers OLIGO320 and OLIGO309, digested with EcoRI + BamHI, and inserted into pEGFP-HIS-BAF (internal Lab ID: PLA1080).

#### 2.4. Plasmids used

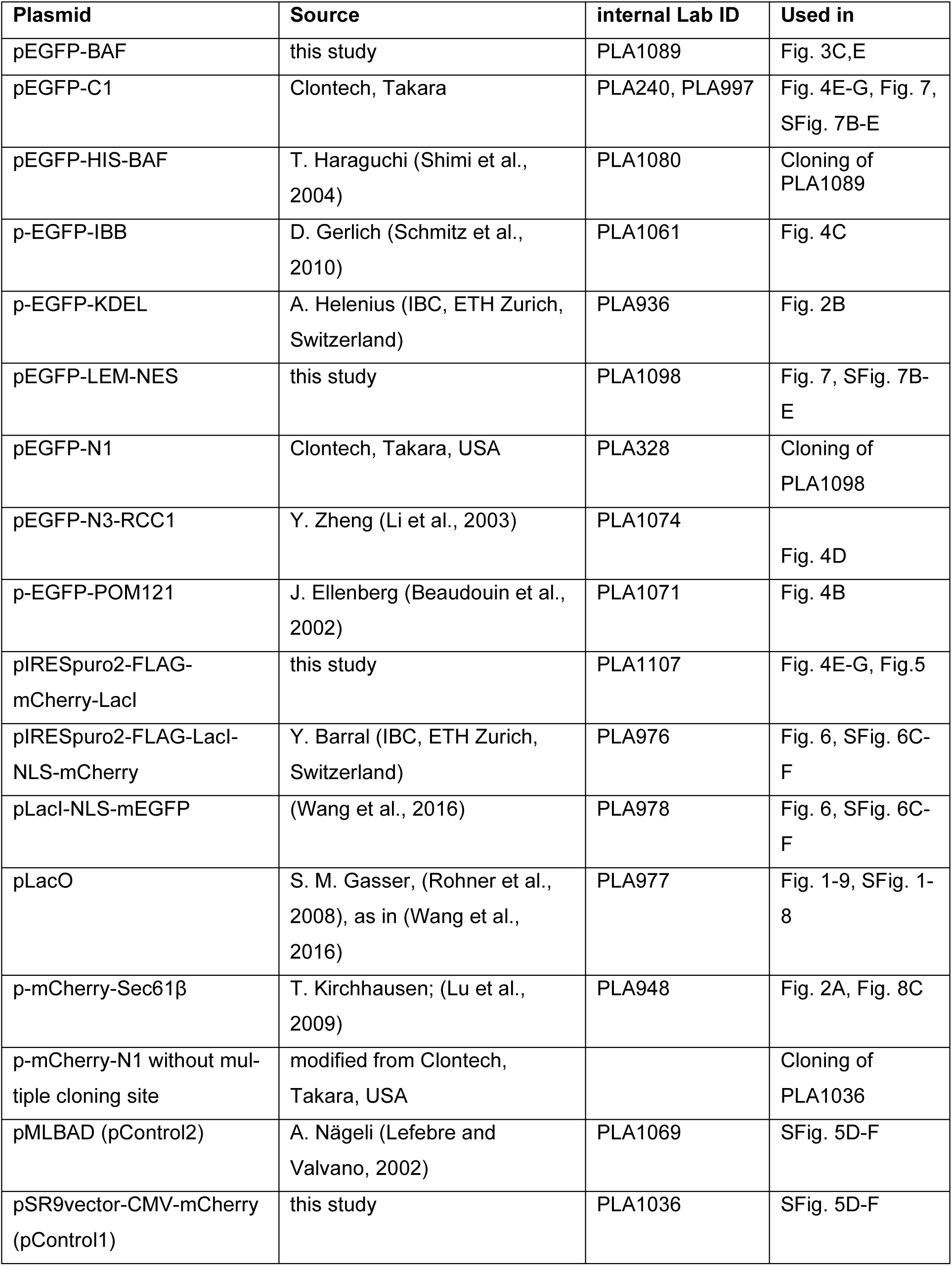

#### 2.5. Plasmid transfection

**Lipofection:** Plasmid was lipofected into cells using X-tremeGENE 9 DNA Transfection Reagent (Roche; Basel, Switzerland). The plasmid:transfection reagent ratio (w:v) was 1:3. Plasmid DNA concentration was either 25 ng (Fig. 2A, B, Fig. 3, Fig. 4, Fig. 5, Fig. 8, Fig. 9, SFig. 8, SFig. 5C HeLa, E, F), 100 ng (Fig. 1, Fig. 2C, Fig. 6A-D, Fig. 7, SFig1, SFig. 2, SFig. 3, SFig. 5, SFig. 6A, SFig. 7), or 330 ng (SFig. 5 MDCK part) per cm^2^ cell culture dish area. For double transfections (Fig. 4E-G, Fig. 6, SFig. 6C-F) plasmids were mixed in a 1: 1 ration and transfected at total 100 ng/cm^2^ cell culture dish area. To wash away excess transfection mix cells were washed with 20 U/ml heparin (Sigma Aldrich, Switzerland) 6 hours after lipofection in Fig. 3B-D, Fig. 4A-D, SFig. 6B. In all other condition, the lipofection mix was left to incubate with the cells for the time mentioned.

**Electroporation:** Electroporation was conducted by a MicroPorator (AxonLab; Baden, Switzerland) with Neon Transfection system 10 µL Kit (invitrogen; Waltham, Massachusetts, United States). The electroporation parameters were 1000 V, 30 ms and 2 pulses for 10 µl electroporation tips using 250 ng DNA per 10^5^ suspension cells in R-buffer. Electroporated cells with same condition were collected in a tube and then seeded on cover slips (SFig. 1B, SFig. 5A, B, Fig. 7, SFig. 7),

### 3. FISH

#### 3.1. FISH probes

##### FISH probes of PNA quality

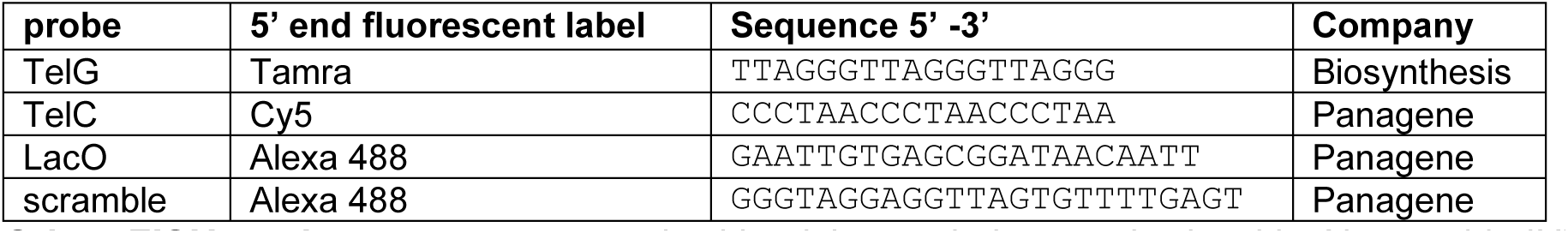

**Other FISH probes** were generated with nick-translation method, with Alexa 568-dUTP (Invitrogen), according to manufacturer’s instructions on indicated template DNAs.

#### 3.2. DNase I enzyme treatment prior to FISH

U2OS and HeLa K cells were fixed with methanol (Supelco; Bellefonte, Pennsylvania, United States) for 10 min at −20 °C and then washed three times in 1x PBS. Cells were permeabilized with 0.5 % Triton X-100 (Sigma-Aldrich, St. Louis, Missouri, United States) for 10 min, then incubated with 0.5 unit/µl DNase (BioConcept; Allschwil, Switzerland) in 1 x PBS for 2 to 2.5 hours at 37 °C.

#### 3.3. Regular FISH

Method modified from (Lansdorp et al., 1996). Cells were rinsed briefly in PBS before fixation. The cells were fixed in 2 % paraformaldehyde (Polyscience; Hirschberg an der Bergstrasse, Germany) in 1x PBS pH 7.4 for 10 min at room temperature (RT) or in 100 % methanol for 10 min at −20 °C. Cells were rinsed in 1x PBS three times for 5 min and fixed again for 10 min in methanol at −20 °C if they were fixed with 2 % PFA before. Cells were permeabilized with 0.2 % Triton X-100 for 10 or 20 min, then treated with PBS containing 20 mg/ml RNase (Thermo Fisher Scientific) at 37 °C for 30 min to 1 h. PNA probes were diluted to 20 nM concentration in hybridization solution (70 % deionized formamide (Eurobio; Paris, France), 0.5 % blocking reagent (Roche), 10 mM Tris-HCl (pH 7.2)). The DNA was denatured at 80 °C for 3 or 15 min. And then incubated in a humid chamber in the dark for 2 hours at RT. Cells were washed with hybridization wash solution 1 (10 mM Tris-HCl (pH 7.2), 70 % formamide and 0.1 % BSA (Gerbu; Gaiberg, Germany)) for two times, 15 min each time at RT and with hybridization wash solution 2 (100 mM Tris-HCl (pH 7.2), 0.15 M NaCl and 0.08 % Tween-20 (Sigma-Aldrich)) for three times. The nuclei were stained by Hoechst33342 (Thermo Fisher Scientific) for 3 min at RT in 1 x PBS and rinsed once with 1x PBS. Cover slips containing cells were mounted in Mowiol 4-88 (Sigma Aldrich) containing 1.4 % w/v DABCO (Sigma Aldrich), sealed with nail polish. This method was applied for the results shown in Fig. 8A, B, SFig. 5E, F, SFig. 8 A-E.

#### 3.4. IF-FISH

After the RNase treatment, cells were blocked with 5 % BSA in 1 x PBST for 1 hour at RT. Then cells were incubated in primary antibody diluted in 1% BSA in 1 x PBST in a humidified chamber for 1 hour at RT. Incubation with the secondary antibody (1:500 for each) in 1 % BSA / 1 x PBST for 1 hour at RT in the dark. Cells were fixed with 4 % paraformaldehyde for 7 min, and then PFA was quenched with 5 % BSA in 1 x PBS and 20 mM glycine for 30 min. Cells were hybridized with probes as described above. This method was applied for the results shown in Fig. 8C, E, Fig. 9, SFig. 8F-H.

### 4. Immunofluorescence

#### 4.1. Antibodies used

| Antibodies | Source | host | fixation | dilution | Identifier | internal Lab ID |
| --- | --- | --- | --- | --- | --- | --- |
| monoclonal NPC (Mab414) | Abcam | mouse | MeOH | 1:1000 (U2OS) or 1:2000 (HeLa) | ab24609 | AB324 |
| Lap2β | BD transduction laboratories | mouse | formaldehyde | 1:500 | 611000; 27/LAP2 | AB273 |
| polyclonal Emerin | Abcam | rabbit | formaldehyde | 1:500 | ab40688 | AB286AB321 |
| serum LEM4 | Ian Mattaj, (Asencio et al., 2012) | rabbit | formaldehyde | 1:1000 | BCFED3 20.1.10 | AB282 |
| polyclonal LBR | abcam | rabbit | MeOH |  | ab122919 | AB264 |
| serum ELYS/MEL-28 | Iain Mattaj (Franz et al., 2007) | rabbit | formaldehyde | 1:200 | N/A | AB304 |
| IgG, Alexa-Fluor™ 647 | ThermoFischer Scientific | mouse | - | 1:500 | A21236 | AB251 |
| IgG, Alexa-Fluor™ 594 | ThermoFischer Scientific | mouse | - | 1:500 | A11032 | AB250 |
| IgG, Alexa-Fluor™ 647 | ThermoFischer Scientific | rabbit | - | 1:500 | A21245 | AB316 |
| IgG, Alexa-Fluor™ 594 | ThermoFischer Scientific | rabbit | - | 1:500 | A11037 | N/A |
| IgG, Alexa-Fluor™ 488 | ThermoFischer Scientific | rabbit | - | 1:500 | A11034 | AB252 |

#### 4.2 Immunofluorescence staining

Cells in Fig. 3B were fixed 30 hours after pLacO transfection. Cells were either fixed with methanol at −20 °C for 6 min, or with 1 % or 4 % paraformaldehyde for 10 min at RT. Cells were permeabilized for 5 min or 10 min with 0.2 % or 0.1 % TritonX-100 at RT. Blocking was performed with 5 % Bovine serum albumin (Boehringer Mannheim, now Roche) in 1x PBST (1x PBS with 0.05 % Tween-20) for 1 hour at RT. Cells were then incubated with primary antibodies diluted in blocking buffer for 1 hour. Followed by incubation with secondary antibodies diluted in blocking buffer for 45 min to 1 hour. Cells were then stained with 2 µM Hoechst33342 (Molecular probes, Thermo Fisher Scientific) for 10 min and mounted in Mowiol with 1.4 % w/v DABCO. Cover slips were sealed with Nail polish (Lucerna Chem AG; Luzern, Switzerland) and stored at 4 °C.

### 5. Image acquisition

Imaging was done at the Scientific Center for Optical and Electron Microscopy (ScopeM, ETH Zurich).

#### 5.1. Fixed cell imaging

For images of fixed cells, z-stacks minimally encompassing entire cells were acquired in 0.3 µm or 0.2 µm steps using a 60x NA 1.42 objective on a DeltaVision personalDV multiplexed system (epifluorescence based IX71 (inverse) microscope; Olympus; Tokio, Japan) equipped with a CoolSNAP HQ camera (Roper Scientific; Planegg, Germany).

Results shown in Fig. 7, SFig. 7 employed imaging using the DeltaVision personalDV multiplexed system with a 60 x 1.42NA DIC Oil PlanApo Objective and a pco.edge 5.5 camera. Z-stacks were acquired with 0.3 µm steps.

A Nikon Wide Field microscope (Nikon Ti2-E; Nikon, Tokio, Japan) was used in Fig. 1A-D, SFig. 1, SFig. 2, SFig. 3 with the S Fluor 20x NA 0.75 DIC N2 WD 1.0mm. For Fig. 6 and SFig. 6F, Plan Apo lambda 60x NA 1.4 oil WD 0.13mm was used. Z-stacks with 41 slices x 0.3 um (12 mm total) were acquired, Dapi (Hoechst, DNA) channel was used as reference and the chromatic offset in mCherry and GFP channels was corrected for.

#### 5.2. Live-cell microscopy

For live-cell microscopy, three cell lines were used: Control HeLa, HeLa-LacI, or HeLa K cells stably expressing aa 244–453 of Lap2β-GFP transiently overexpressing LacI-mcherry. For results displayed in Fig. 1E, F, Fig. 2A, B, Fig. 3C, Fig. 5, SFig. 4 cells were seeded on Lab-Tek II chambers (Nunc, Thermo Scientific) with CO_2_-independent media (Gibco) containing 10 % FCS and incubated at a 37 °C on a Spinning Disk microscope (Nipkow spinning disk setup with Nikon Eclipse T1 (inverse) microscope, equipped with 2x EMCCD Andor iXon Ultra cameras, LUDL BioPrecision2 stage with Piezo Focus, Carl Zeiss Microscopy; Jena, Germany). For imaging in Fig. 5 the GFP-Like (Em 520/35) and DsRed-like (Em 617/73) Emission Filter Wheels and 2x Evolve 512 cameras (Photometrics; Tucson, Arizona, United States) were used. For long-term time-lapse imaging (Fig. 5, SFig. 4), cells were recorded every 15 min in z-stacks (33x 0.7 μm steps using a 63x 1.2 NA objective). To monitor cell contours, cells were illuminated with transmission light with single z-focus. For some still images cells expressing Sec61-mCherry (Fig. 2A), eGFP-KDEL (Fig. 2B) and eGFP-BAF (Fig. 3C) were imaged after incubation with 2 µM Hoechst33342 for 10 min, using a DeltaVision microscope (DeltaVision personalDV system (epifluorescence based IX71 (inverse) microscope; Olympus).

For results displayed in Fig. 1, SFig. 2, SFig. 3, HeLa Control cells were seeded on ibidi 8-well chambers (ibidi µ-Slide 8 well ibiTreat, Gräfelfing, Germany). 24 hours after seeding, cells were incubated at 37 °C with 5% CO_2_ (OkoLab, Pozzouli NA, Italy) either at the Visitron Spinning Disk (experiments e1 (internal Lab ID: EXP345) and e2 (internal Lab ID: EXP337)) or a Nikon Wide Field microscope (Nikon Ti2-E (inverse), experiments e3 and e4 (internal Lab ID: EXP604)). For Visitron spinning disk imaging a GFP-Like (Em 520/35) Emission Filter Wheel and 2x Evolve 512 cameras (Photometrics) were used. Bright field imaging was done with the coolLED *p*E-100 control system (coolLED; Andover, Great Britain). For Nikon Wide Field imaging the GFP (Em 515/30) Emission Filter Wheel or bright field pre-setting was used. For detection, the Orca Fusion BT (Hamamatsu; Shizuoka, Japan) (2304×2304 pixels, 6.5 µm x 6.5 µm) system was used. For each experiment at the Visitron Spinning Disk microscope, 5 regions of interest (ROI) were imaged. For each experiment at the Nikon Wide Field microscope, 6 ROI were imaged. In the live-cell analysis included are only ROI with 0.9766 cells/pixel, thus one ROI of e1 at the Visitron Spinning Disk microscope and two ROI of e3 as well as two ROI of e4 at the Nikon Wide Field microscopewere excluded. Cells were recorded every 30 min as z-stacks (22 x 0.7 μm steps using 20x 0.75 CFI Plan Apo VC at the Visitron Spinning disk and 22 x 0.7 μm steps using S Fluor 20 x NA 0.75 DIC N2 WD 1.0mm at the Nikon Wide Field). On both microscopes, cells were lipofected with pLacO using X-tremeGENE 9 DNA Transfection Reagent (Roche). The plasmid:transfection reagent ratio (w:v) was 1:3 with a plasmid DNA concentration of 100 ng/cm^2^. The lipofection mix remained on the cells during imaging. The death rate was low at both microscopes (SFig. 2A).

#### 5.3. FRAP

FRAP experiments (Fig. 4E-G) were performed using a modified method that was previously reported (Clay et al., 2014). 24 hours after pLacO transfection, live HeLa-LacI cells (seeded on a Lab-TekTM II chamber, CO_2_-independent media, 37 °C incubator) and free eGFP were imaged on a confocal microscope (LSM 760; Carl Zeiss Microscopy) with a Plan Apochromat 63x /1.4 NA oil immersion objective. The ZEN software (Carl Zeiss Microscopy) was used to control the microscope. eGFP emission was detected with a 505 nm long pass filter. Photobleaching was applied on a region of interest (cluster and then control area) as indicated in Fig. 4E-G. Bleaching was applied with 50-100 iterations using 30-50 % laser power, but always with the same settings between the cluster and control area in each cell.

#### 5.4. Correlative light and electron microscopy (CLEM)

HeLa K cells stably expressing LacI-NLS-mCherry were cultured on a 3.5 cm glass bottom dish with grid (MatTek; Ashland, Massachusetts, United States) and transfected with pLacO for 24 hours. Cells were processed as described in (Wang et al., 2016).

### 6. Data Analysis (Fiji, Prism, Diatrack, etc.)

#### 6.1. Image processing

Images acquired from DeltaVision (Olympus) microscope (Fig. 2C, Fig. 3B-D, Fig. 4A-D, Fig. 8, SFig. 9A, SFig. 5A, E, SFig. 6A, SFig. 7C, D, SFig. 8A, B, D) were deconvolved using Softworx (Applied Precision; Rača, Slowakia). Images acquired from LSM 710 confocal microscope were deconvolved using Huygens Software (Scientific Volume Imaging; Hilversum, Netherlands) before correlating with EM images (Fig. 2A). The correlation analysis between confocal and EM images (Fig. 3A) were performed using Amira software (FEI/Thermo Fisher Scientific), as in (Wang et al., 2016). General, the presented images are single z-slices or if indicated projections of multiple z-slices images.

Images in Fig. 3A (confocal image), Fig. 6 and SFig. 6F were deconvolved using Huygens (Scientific Volume Imaging).

#### 6.2. Image analyses

Images were analyzed using Fiji 1.51n Software.

##### 6.2.1 Colocalization

For co-localization analyses, the overlay of the reporter fluorescence and LacI fluorescence was used (Fig. 2C, Fig. 6, Fig. 8, SFig. 6D, E, SFig. 8). The qualitative classes for reporter molecules “enriched”, “non-enriched”, “present” and “absent” are established applying the following rules:

For experiments presented in Fig. 3, Fig. 4:

Generally, marker fluorescence intensities were used to qualitatively determine co-localization of markers with the plasmid focus with the following criteria.

“Non-enriched”: plasmid foci with marker fluorescence signal in the z-stack slice in, directly underneath or above the position of the plasmid focus, and/or marker fluorescence signal in xy-direction adjacent. The intensity of the marker fluorescence is like the cytoplasmic marker fluorescence (relative readout to the intensity of the rest of the cell).

Plasmid foci with marker “enriched” have marker fluorescence at same positions described in “non-enriched”, but with higher intensities compared to the reference marker fluorescence of the respective marker (i.e., Emerin at NE or in ER; LEM4 in ER). Where sensible, results state the two “enriched” reference marker fluorescence (i.e., ER or NE).

Category “present” (Fig. 8 and Fig. 4) encompasses “enriched” and “non-enriched”.

Plasmid foci with marker “absent” do not have marker fluorescence in the adjacent slides, underneath or above the position of the focus, nor a marker fluorescence signal in xy direction adjacent to the focus nor in the sliced with the focus.

For data presented in Fig. 3E:

For Emerin and Lap2β, the quantitative enrichment factor analysis (below) were back translated into qualitative classification: "enriched” with an enrichment factor >1, or “non-enriched” with 0>enrichment factor>1, or “absent” for enrichment factor being zero. For BAF and LBR, the classification of “enriched”, “non-enriched”, and “absent” as described above for Fig. 3 was used.

For data presented in Fig. 6, SFig. 6C-F:

In the single z-slice, where the plasmid focus was in focus, a line scan across the biggest axis of the plasmid focus and either the ER (for LEM4) or across the nucleus (for Emerin) was made, displaying the intensity distribution along that line (Fiji, line scan). Classification was according to the following intensity criteria: enriched > NE or > ER: the average intensity of the reporter (Emerin or LEM4) at the plasmid focus was higher compared to the average reporter intensity at the NE or surrounding ER, displayed along the line. Like ER: fluorescence of the reporter (Emerin or LEM4) was in average identical the intensity in the ER surrounding the plasmid focus. Absent: no intensity of the reporter (Emerin or LEM4) at the plasmid focus.

For the experiments presented in Fig. 7:

Intensities of reporter proteins (LEM4 or Emerin) at the plasmid focus were visually compared to intensities reporter proteins in the surrounding cytoplasm. “Present”: If the reporter intensity was equal or higher at the plasmid focus than that of the surrounding cytoplasm in the focal slice, directly underneath or above the focus-position of the plasmid focus (0.3 µm distances). Otherwise, the classification was “absent”.

For the experiments presented in SFig. 7B, C:

The intensity of Emerin immunofluorescence was measured as RawIntDen (Fiji) in a square (20×20 px) covering ER and in a same sized square covering the nucleus of a single cells in maximum intensity projected images. Chosen were in both cases regions where the intensity appeared the most intense as judge by eye. The ratio between the RawIntDen value at the NE divided by that at the ER was calculated for each cell with minimally 1 cytoplasmic plasmid focus. A ratio above 1 reports about a higher intensity of Emerin (and therefore more Emerin) at the NE compared to the ER of that same cell. A ratio below 1 represents a higher intensity of Emerin (and therefore more Emerin) at the ER compared to the NE of the same cell.

Quantitative enrichment factor analysis (SFig. 6B):

Single z-slice images were analyzed. The fluorescence intensity was measured along a line crossing the plasmid focus and the nucleus. Along this line, the fluorescent intensities of two brightest pixels at the edges of the plasmid focus (I (c1), I (c2)) or the nucleus (I (n1), I (n2)), were averaged. Another averaged intensity of 30-50 pixels along this line, in a cytoplasmic region, was used as background intensity (I (background)). The enrichment factor was calculated as: (Enrichment factor = ((I(c1)+I (c2))*0.5 - I (background))/ ((I (n1)+I (n2))*0.5 - I (background)).

##### 6.2.2 Live-cell imaging analyses

**Plasmid focus localization** (Fig. 1, SFig. 2, SFig. 3):

For plasmid focus analyses, two interphase localizations are classified: cytoplasmic and nuclear. <u>Cytoplasmic plasmid foci</u>: These intracellular LacI-positive plasmid foci are outside of the volume marked by LacI-NLS-GFP fluorescence reporting about the nucleus or are at the cytoplasmic side of the nuclear envelope, which is reported by the outer boarder of the nuclear LacI-NLS-GFP fluorescence, but with minimally 1 pixel of background intensity between the LacI intensity at the plasmid focus and that of the nucleus. <u>Plasmid foci in the nucleus</u> are either nucleoplasmic LacI-positive foci or LacI-positive foci at the nucleoplasmic side of the nuclear envelope, which is reported by the outer boarder of nuclear LacI-NLS-GFP fluorescence. Nucleoplasmic plasmid foci (nuclear foci) are defined as plasmid foci inside the volume of nuclear LacI-NLS-GFP fluorescence but with an intensity higher than that of the general nuclear LacI-NLS-GFP fluorescence. In addition, the z-slice in which the plasmid focus is most in focus (focal-z-slice) and at highest intensity is also the z-slice, in which nuclear LacI-NLS-GFP fluorescence covers the biggest area. Further, the focal-z-slice of the plasmid lies in between other z-slices in which the general nuclear LacI-NLS-GFP fluorescence is still in focus. Typically, depending on the z-stack spacing, the general nuclear LacI-NLS-GFP fluorescence is still in focus within +/- 1 µm of the focal plane of the plasmid focus. This is to be compared to a situation, where a plasmid focus is at the nuclear envelope and thus in an upper z-slice. In this case the plasmid focal plane is not identical with the z-slice of the biggest area of general nuclear fluorescence. These classification criteria were used in Fig. 1, SFig. 2, SFig. 3 as well as for classification of plasmid foci being formed in the nucleoplasm in SFig. 5G+H. We chose these strict conditions to exclude the option of a false positive nuclear assignment to plasmid foci.

**Origin-destination analysis** (Fig. 1D, SFig. 2H):

To avoid analyzing plasmid foci that formed in the cytoplasm but in close proximity to the NE, we excluded plasmid foci that formed at the inner side of the nuclear periphery from the analysis focusing thus on foci formed in the inner nucleoplasm. To allow for sorting time the last 25 % of the forming plasmid foci in the pooled data set were excluded from this analysis. For the “origin-destination” analysis the location of formation of each plasmid focus was noted (“origin”) (either interphase: cytoplasm or nucleoplasm, or during mitosis). Then, after tracing the focus over time until the end of imaging, the location of each plasmid focus at imaging end (“destination”) was noted. If during imaging a cell fused with another cell, died, produced a micronucleus, the nucleus fragmented, the plasmid focus disappeared, or the cell was in mitosis at the end of imaging the focus localization in the last frame before one of these events was noted as the corresponding destination of the given plasmid focus. Only cells that completed a mitosis (first frame with two distinct nuclei in LacI-NLS-GFP channel and two fully divided cells in brightfield channel) were analyzed.

**FRAP quantification** (Fig. 4E, F):

Using Fiji, the mean fluorescence recovery signal was quantified in the bleached area. The fluorescence signal was normalized to that at the beginning of the experiment. All experiments were transferred to Prism software (GraphPad) and fit on an exponential FRAP curve, the mobile fraction was measured by determining the half time (t _1/2_) of fluorescence recovery to reach a plateau level.

**FISH co-localization analysis** (Fig. 8, Fig. 9, SFig. 8):

Fluorescence signals of stained proteins were boosted until the background level of the cell’s cytoplasm was visible. Proteins co-localized at telomeric DNA if the signals at the telomeric DNA foci were visually than the background in close vicinity and the boosted setting.

### 7. Statistics

Statistics were conducted using Prism 8.0.0 (GraphPad) built-in analysis tools, methods used are indicated in the figure legends.

Data were tested with Gaussian distribution for normality (*D’Agostino & Pearson* normality test) (α=0.05)) if t-tests were used.

## Acknowledgements

We thank Yves Barral (ETH-Zürich), for continued discussions and support of this study. We thank Madhav Jagannathan, members of the Kroschewski and Barral labs for helpful discussions and comments on the manuscript. We acknowledge financial support by Swiss National Science Foundation (grant 31003A_172908/1 to RK) and the Novartis foundation for medical-biological research (grant 20C229 to RK).

---

**SFig. 1.**
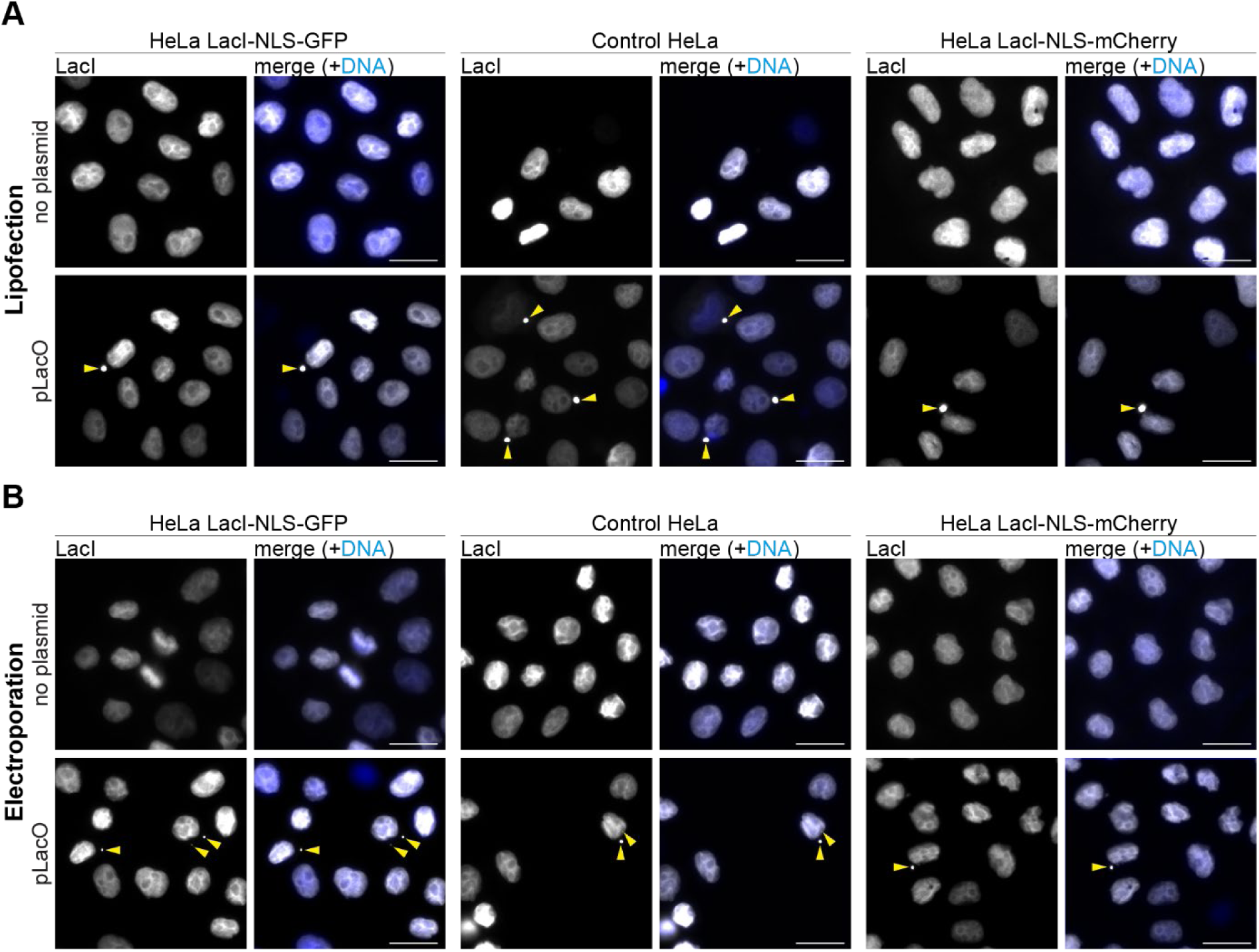
Only plasmid-transfected cells have cytoplasmic LacI foci. (**A, B**) Images of HeLa-LacI cells stably expressing LacI-NLS-GFP or LacI-NLS-mCherry lipofected (**A**) or electroporated (**B**) without or with pLacO and fixed 24 hours after transfection. Scale bar, 20 µm. DNA, blue (Hoechst stain). Plasmid foci in the cytoplasm, yellow arrowheads.

**SFig. 2.**
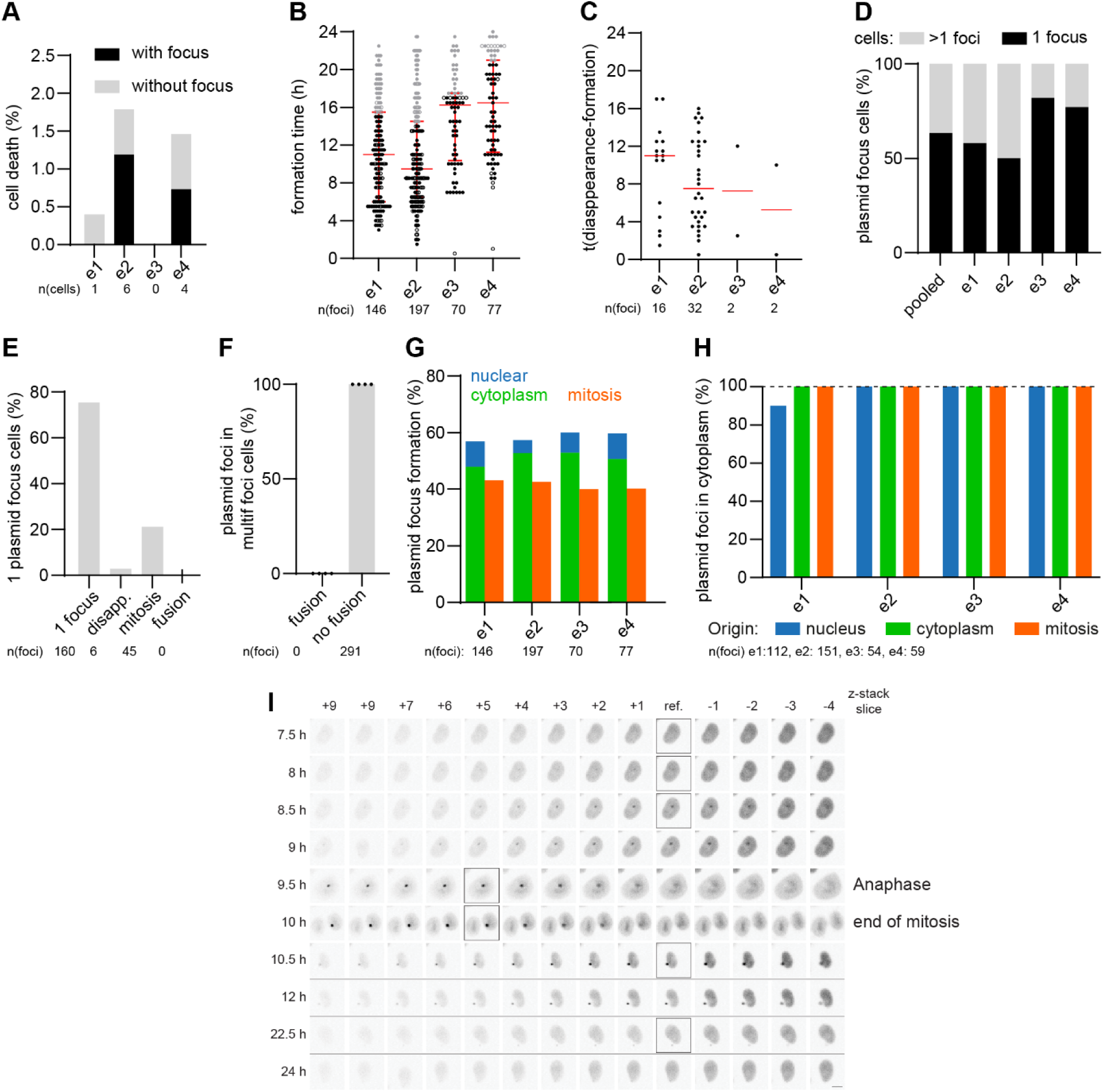
Dynamics of plasmid foci in individual live cell imaging experiments. Individual exp., e1 - e4. (**A**) Cell death events until end of imaging. Cells died without focus, grey; died with focus, black; % relative to cells at end. n(cells): 253, 336, 270, 273. (**B**) Timing of focus formation. 1 focus, circle; time, after lipofection; median & interquartile range, red; last 25 % formations, grey. (**C**) Presence period of disappearing foci. 1 focus, circle; median, red line. (**D**) Cumulative frequency of 1-focus cells (black) and multi-foci cells (grey) during imaging period. Maximal number of foci per cell during lifetime of single cells; pooled data, pooled. (**E**) Analysis if origin of 1-focus cells at imaging end. Cell formed 1 focus or divided propagating it, 1 focus; cell formed multiple foci and all but one disappeared, disapp.; partitioning in mitosis resulted in 1-focus cell(s), mitosis; foci fused, fusion. Pooled data. n(1-focus cells): 211. (**F**) Fusion of foci in multi-foci cells. Pooled data. (G) Origin of forming foci. Normalized to all foci formed per exp. (**H**) Cytoplasmic foci depending on origin. Last 25 % formations excluded. n(foci): 112; 100 % reference, dashed line. **(I**) Focus formation in the nucleoplasm. Cell as Fig. 1A (nucleoplasmic formation, corresponding images with black squares) with z-slices above and below reference slice (ref.) Scale bar, 10 µm. Time, after lipofection. Black horizontal lines, skipped time points.

**SFig. 3.**
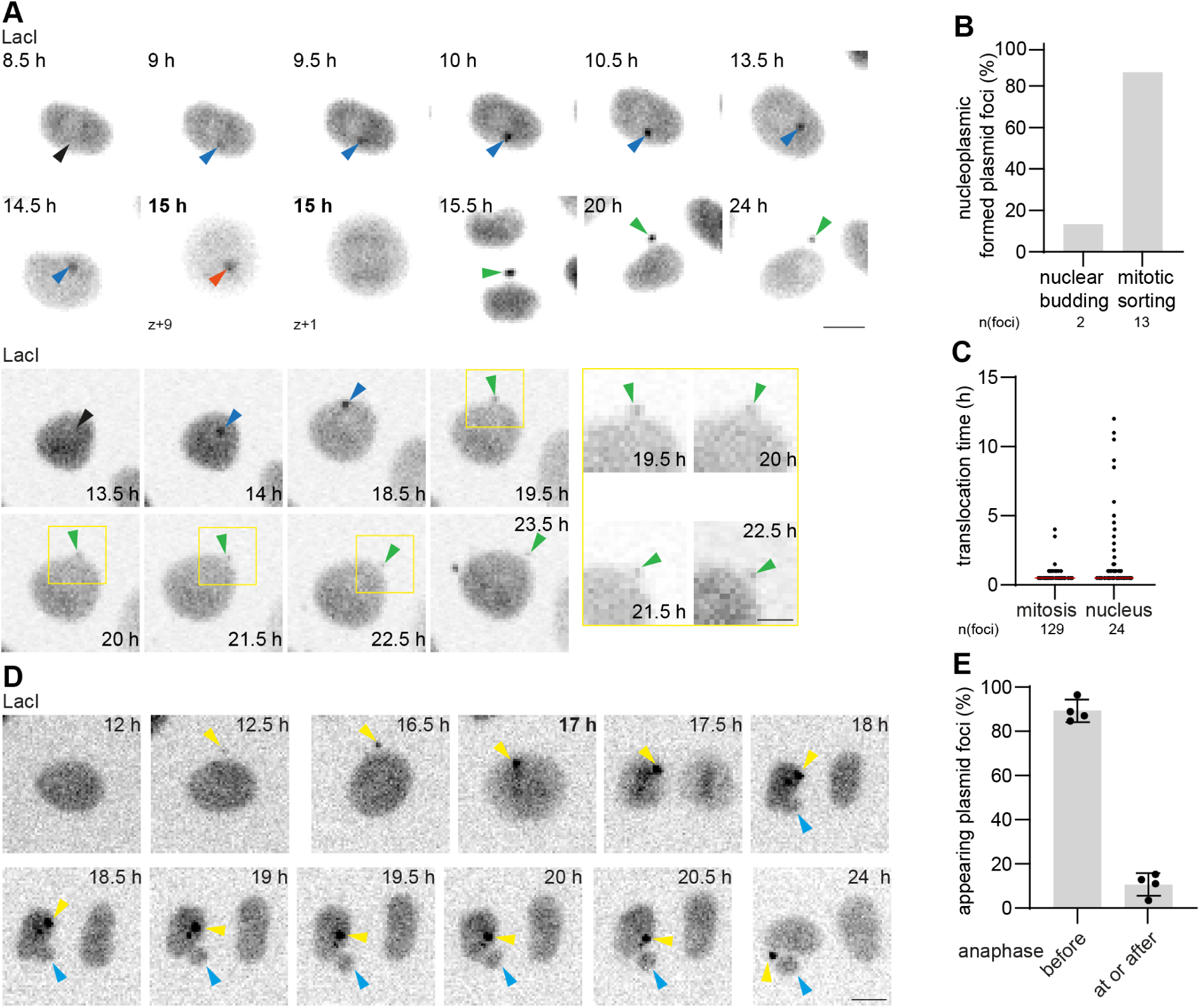
How plasmid DNA leaves the nucleus. (**A**) Two example time-lapse images of focus formations in HeLa-LacI cells; mitotic sorting (upper) and nuclear budding (lower). Scale bar, 10 µm. Time, after lipofection; bold time, mitosis. Arrowheads: nucleoplasmic, blue; cytoplasmic, green; mitosis, orange; future focus formation area, black; single z-slices. (**B**) Quantification of events shown in (**A**). Pooled data of 4 exp.; only foci analyzed, which appeared in the nucleoplasm but not at the border of the nuclear LacI-NLS-GFP fluorescence neither on nucleoplasmic side (see method); n(foci): 15. (**C**) Duration between the first detection of a focus and its first localization in the cytoplasm. Only plasmid foci formed either during mitosis or in the nucleus, appearing during the first 75 % of the formation time of all plasmid foci formations and translocating into the cytoplasm, were analyzed. 1 plasmid focus, circle; pooled data of 4 exp.; median, red line; n(foci): 24. (**D**) Time-lapse images contrasting focus (yellow arrowhead) and mitotic micronucleus formations (light blue arrowhead) in HeLa-LacI cells. Scale bar, 10 µm. Time, after lipofection. (**E**) Plasmid foci formed during mitosis relative to anaphase. 4 exp. (circles); mean & SD; n(foci): 207.

**SFig. 4.**
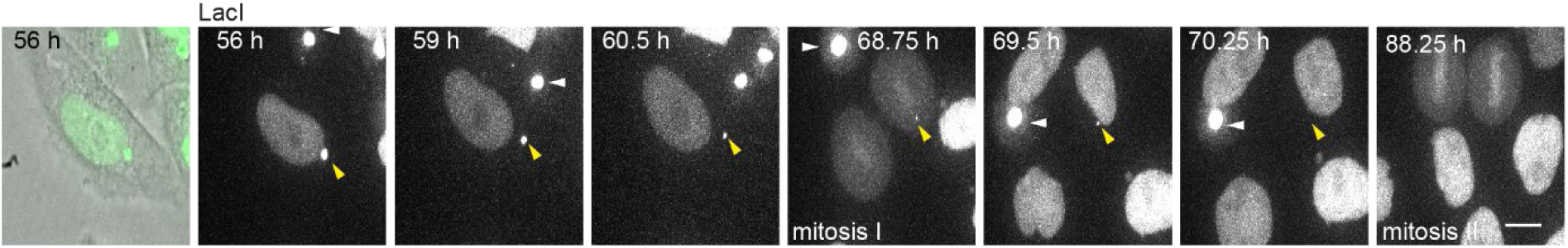
Fluorescence dynamics of plasmid foci in long-term movies. Representative time-lapse images of HeLa-LacI cells showing the disappearance of a focus between 56 hours and 88.25 hours after transfection. Plasmid focus, yellow arrowheads. Persistent brightness of a focus (white arrowhead) in a neighboring cell shows that the disappearance of fluorescence at a focus is not because of bleaching. Cell outline is shown by transmission light in the first frame (56 hours). Scale bar, 10 µm.

**SFig. 5.**
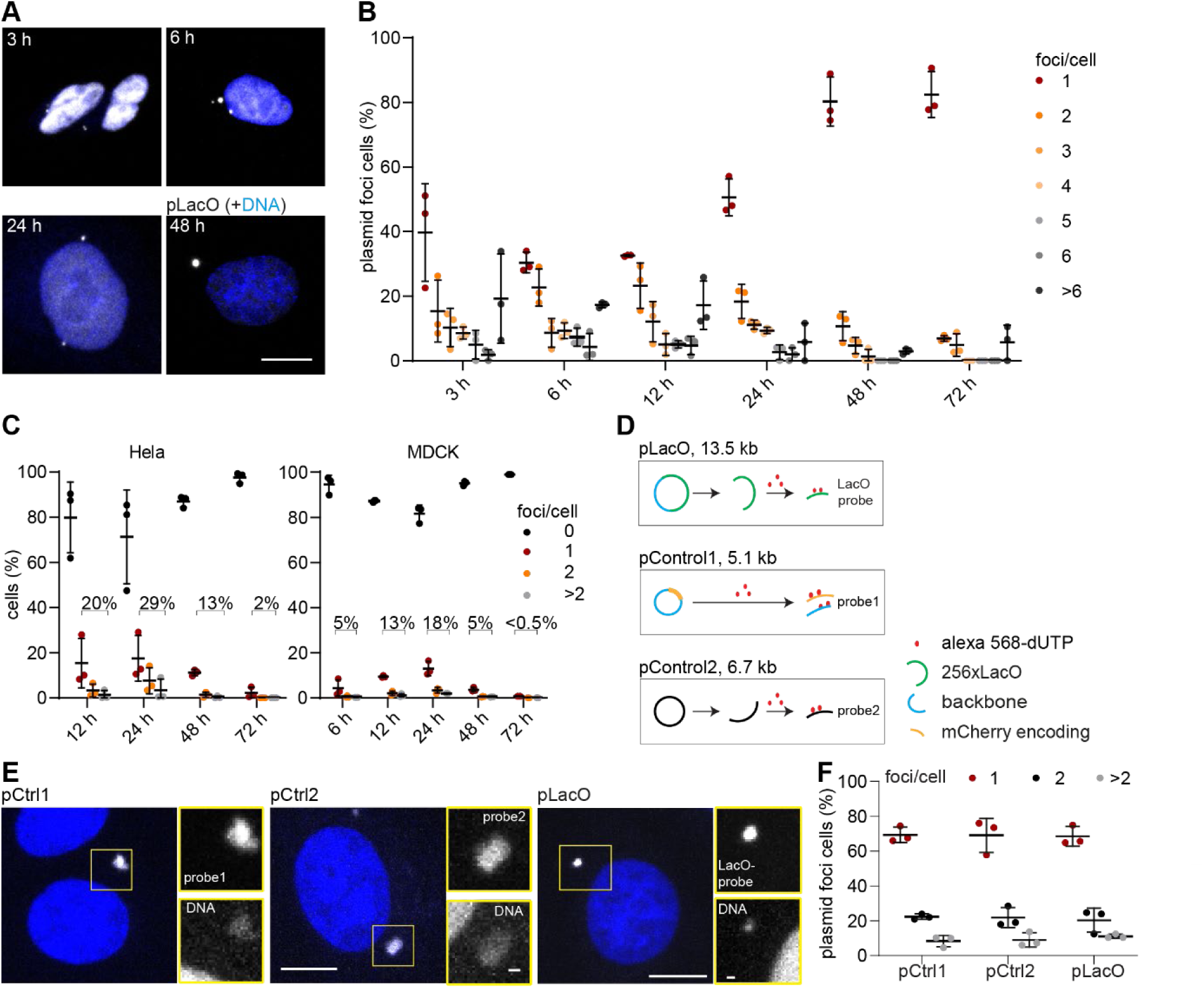
Time, transfection method, and plasmid type dependent reaction towards transfected plasmid DNA. (**A**, **B**) MDCK-LacI cells electroporated with pLacO fixed, imaged, and analyzed at indicated times after electroporation. (**A**) Example images. Scale bar, 10 µm. (**B**) 7 classes (1 - >6) of plasmid foci per cell. 3 exp. 1 exp, circle. Mean & SD; n(cells, 3 hours): 166, n(cells, 6 hours): 162, n(cells, 12 hours): 175, n(cells, 24 hours): 161, n(cells, 48 hours):167, n(cells, 72 hours): 115. (**C**) 4 classes (0 - >2) of plasmid foci per HeLa-LacI (left panel) and MDCK-LacI (right panel) cell at indicated times after lipofection. 3 exp. 1exp., circle; Mean & SD. HeLa-LacI: n(cells, 12 hours): 1092; n(cells, 24 hours): 778; n(cells, 48 hours): 1442; n(cells, 72 hours): 7164. MDCK-LacI: n(cells, 6 hours): 3134; n(cells, 12 hours): 1367; n(cells, 24 hours): 1099; n(cells, 48 hours): 4176; n(cells, 72 hours): 17476.(**D**) Scheme illustrating three transfected plasmids and corresponding FISH probes used in (E, F). (**E**) Representative images of FISH on HeLa-LacI cells lipofected with either pLacO (LacO probe), pControl1 (pCtrl1, probe1), or pControl2 (pCtrl2, probe2) 24 hours after transfection. Images are max. intensity-projected z-stacks. Insets: plasmid foci; scale bars: big images: 10 µm; insets, 1 µm. (**F**) 3 classes (1 - >2) of plasmid foci per HeLa-LacI cell depending on the transfected plasmid. 3 exp, n>50 per exp.

**SFig. 6.**
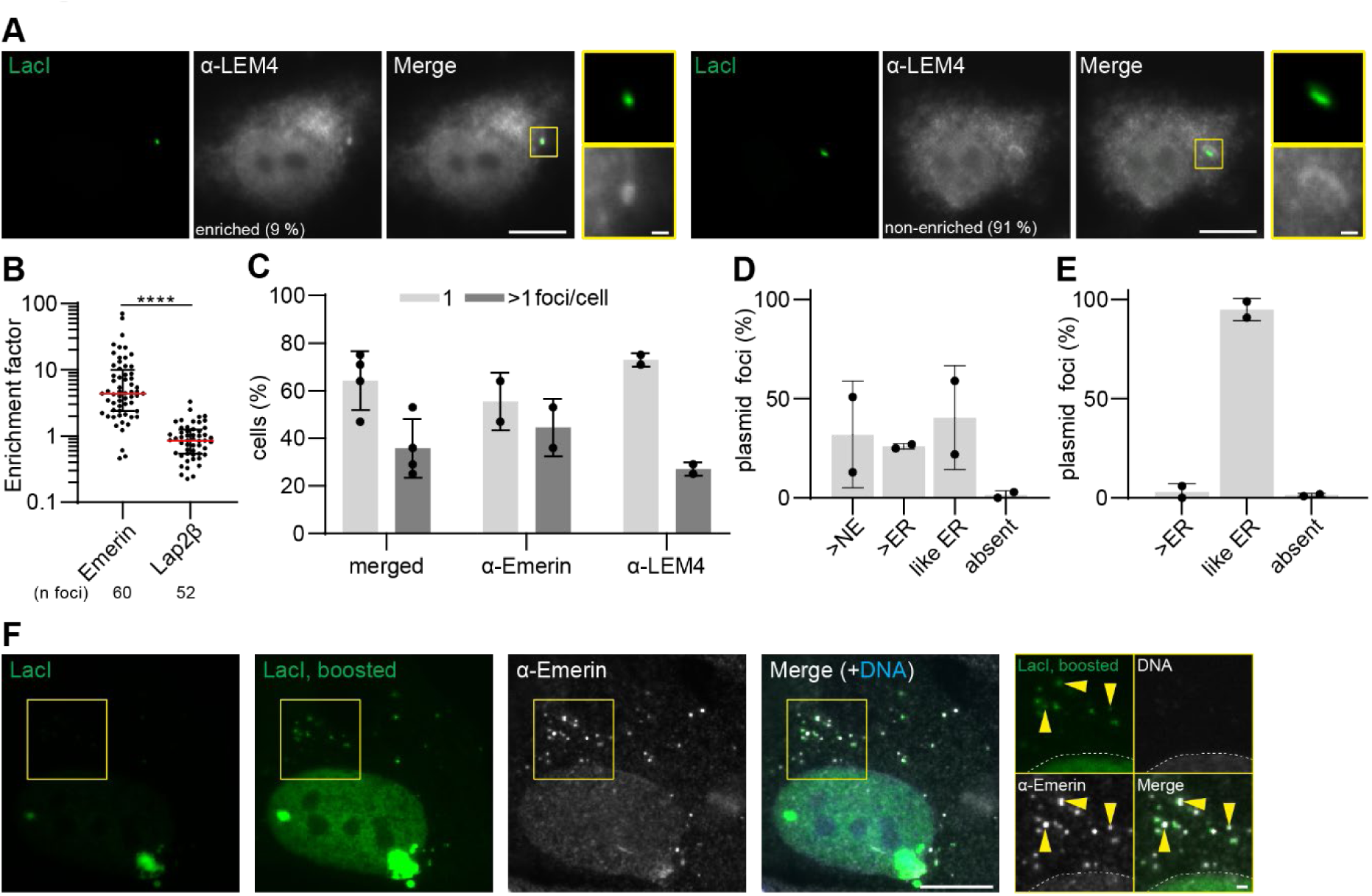
Detailed characterization of the exclusome. (**A**) HeLa-LacI cells electroporated with pLacO and 24 hours later immunostained for LEM4. Single z-slice images; insets: plasmid foci; scale bars: big images,10 µm; insets, 1 µm. Pooled data of 2 exp.; n(cells): 124. (**B**) A ratio-based fluorescence enrichment analysis for Emerin and Lap2β at plasmid foci in HeLa-LacI cells, 24 hours after pLacO transfection. 3 exp. pooled; plasmid focus, circle. Non-paired t-test with log (value); ****: p<0.0001. (**C-F**) Primary human fibroblasts 48 hours after co-lipofection with pLacO and plasmid encoding LacI-NLS-GFP. (**C**) Frequency of 1-focus cells and multi-focus cells. 4 exp. 1 exp, circle; n(cells): 106; mean & SD. (**D, E**) Enrichment classes relative to the NE and the ER based on measured intensities for Emerin (**D**) and LEM4 (**E**) in multi-foci cells. 2 exp., 1 exp., circle; mean & SD; n(Emerin, foci): 227; n(Emerin, cells): 54; n(LEM4, foci): 158; n(LEM, cells): 52. (**F**) Example image of primary human fibroblast 48 hours after transfection and immunostained for Emerin. Single z-slice images. Insets: several plasmid foci; scale bars: in big images 10 µm; in insets: 1 µm; DNA in overview, blue; inset, gray (Hoechst stain). Plasmid foci enriched for Emerin compared to nuclear envelope of same cell, yellow arrowheads. Nucleus outline, dashed line.

**SFig. 7.**
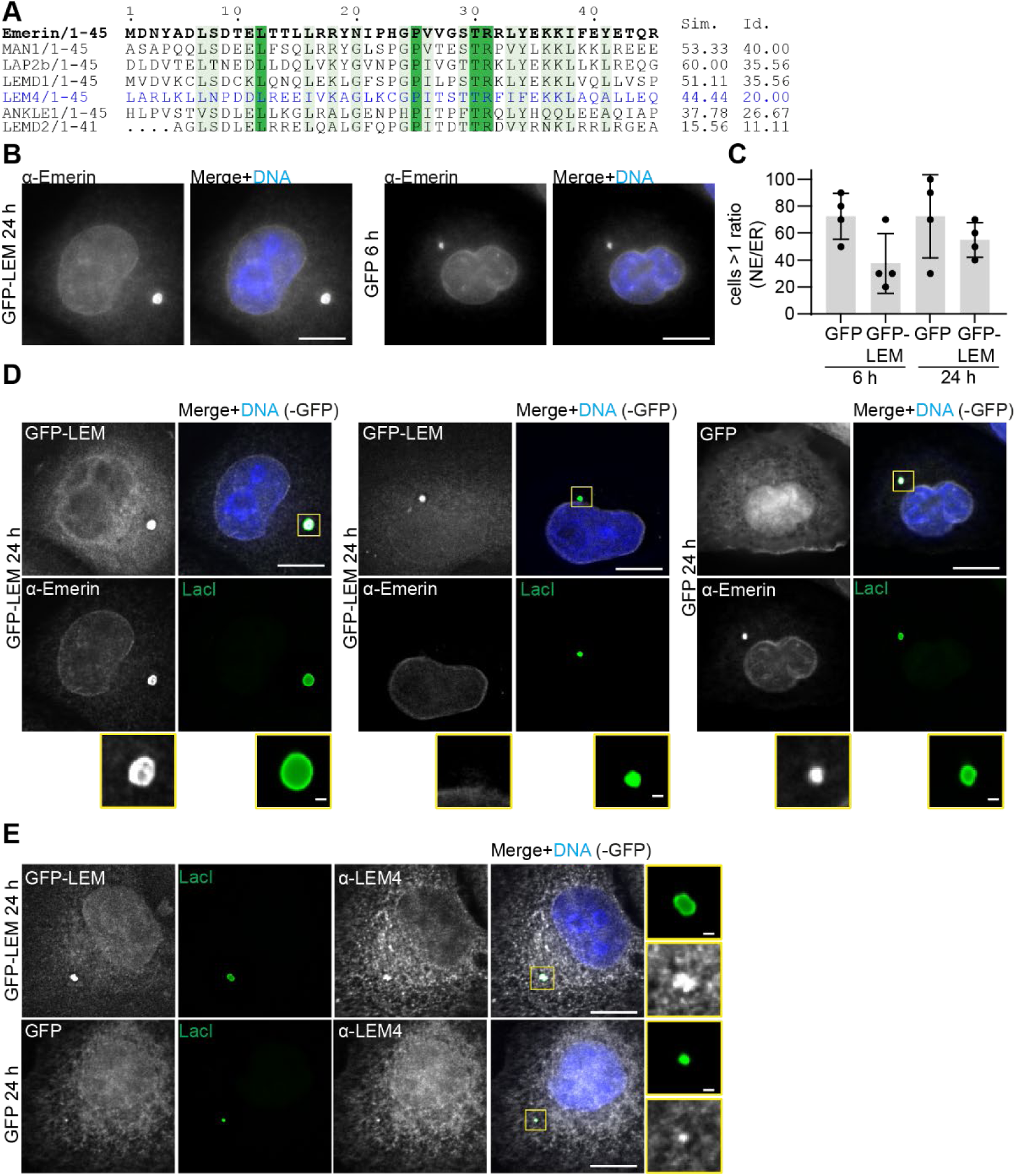
Effects of overexpression of Emerin’s LEM domain. (**A**) Amino-acid alignment for the LEM-domain of various human LEM-domain proteins. Percentage of similar (Sim.) and identical (Id.) residues compared to Emerin (bold). LEM-domain of LEM4, blue. Sim. residues, light green; Id. residues, dark green. (**B-E**) HeLa-LacI cells transiently expressing GFP-LEM or GFP 6 hours (**C**) and 24 hours (**B-E**) after electroporation with pLacO. DNA, blue (Hoechst staining). (**B**) Boosted single z-slice images to visualize ER. Same cells as in (D) (left & right panel) immunostained for Emerin. Scale bar, 10 µm. (**C**) Cells, with a higher measured intensity of Emerin at the NE compared to the ER. 4 exp. 1 exp, circle; mean & SD. (**D, E**) Deconvolved single z-slice images to visualize plasmid foci. Cells were immunostained for Emerin (**D**) or LEM4 (**E**). Insets: plasmid focus. Scale bar: in big images, 10 µm; insets, 1 µm.

**SFig. 8.**
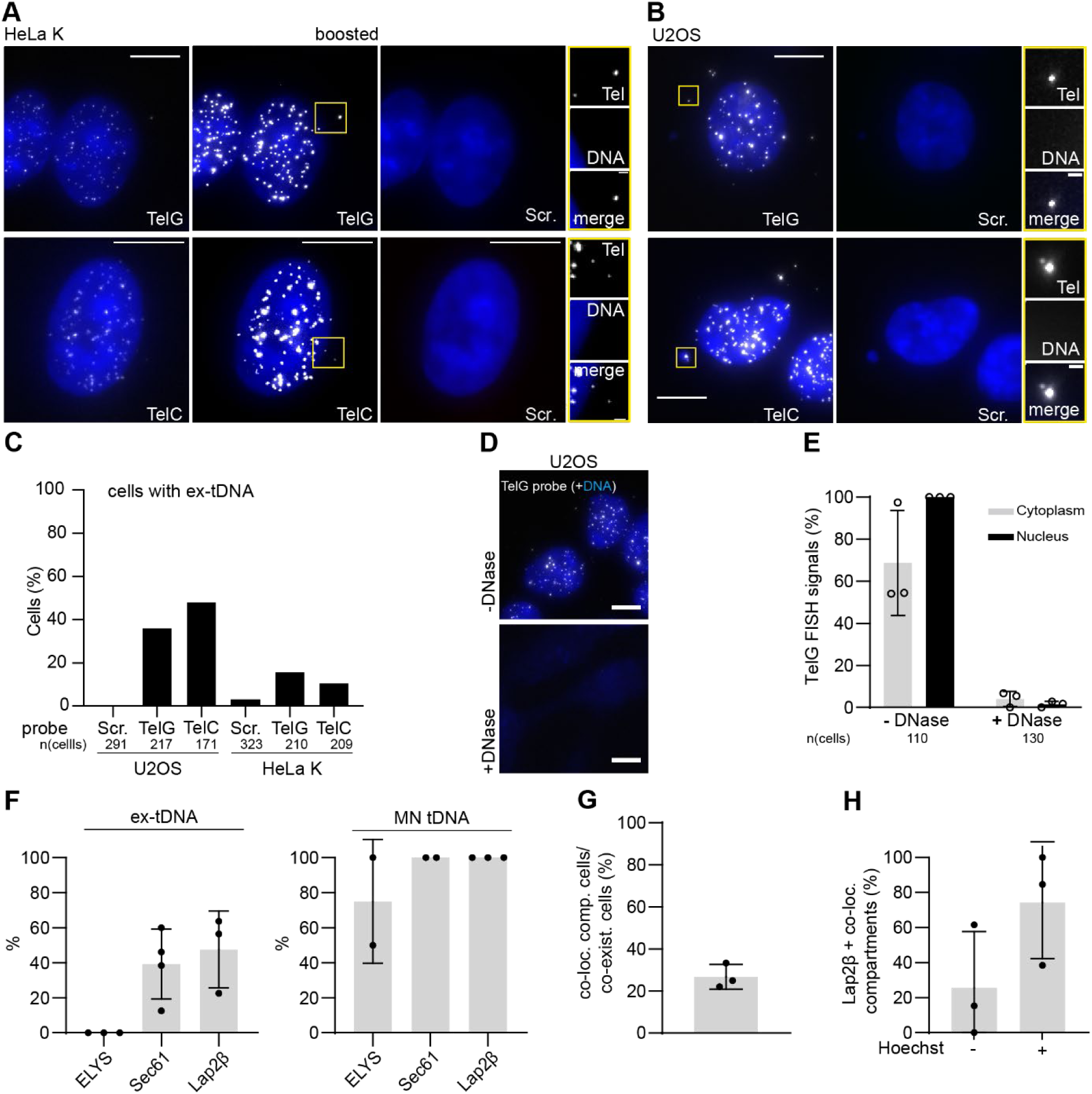
Interphase U2OS and HeLa K cells with ex-tDNA. (**A, B**) Representative max. projected images of HeLa K (**A**) and U2OS (**B**) cells with either TelG or TelC probes plus scramble probes (scr.). Insets: ex-tDNA foci. Imaging conditions of (**A**) and (**B**) were the same; corresponding display; except for when FISH signals were boosted (boosted). Scale bar, 10 µm. (**C**) Percentages of HeLa K, U2OS cells containing ex-tDNA. Pooled data of 3 to 5 exp. U2OS with TelG: 3 exp; n(cells): 47, 73, 97; U2OS with TelC: 3 exp; n(cells): 57, 49, 65; U2OS with scr.: 5 exp; n(cells): 57, 49, 65, 73, 47. HeLa with TelG: 3 exp; n(cells): 55, 59, 96; HeLa-TelC: 3 exp; n(cells): 82, 64, 63; HeLa with scr.: 5 exp; n(cells): 82, 64, 63, 55, 59. (**D, E**) Representative max. projected images (**D**) and quantification of U2OS (**E**) with nuclear and cytoplasmic TelG FISH signals, with and without DNase I treatment. DNA, blue (Hoechst stain). Scale bar, 10 µm; mean & SD; 3 exp. (circles). (**F**) Colocalization of indicated proteins with ex-tDNA (left side) and MN tDNA (right side). 3 exp. (circles); mean & SD; foci: n(Lap2β, ex-tDNA): 154; n(Lap2β, MN tDNA): 12; n(Sec61, ex-tDNA): 95; n(Sec61, MN tDNA): 6; n(ELYS, ex-tDNA): 66; n(ELYS, MN tDNA): 6. (**G**) U2OS cells with minimally one co-localization (co-loc.) compartment (comp.) in cells, which contain both ex-tDNA and plasmid DNA (termed co-existence cells (co-exist.)) 24 hours after pLacO transfection. 1 exp, circle; n(co-exist. cells): 155; n(co-loc comp): 46; mean & SD. (**H**) Quantification of Hoechst signal at Lap2β^+^ co-localization (co-loc.) compartments 24 hours after pLacO transfection. 3 exp. (circles); n(co-existence cells): 46; n(Lap2β+ co-localization compartments): 30; mean & SD.

